# Transformation and Integration of Microenvironment Microarray Data Improves Discovery of Latent Effects

**DOI:** 10.1101/627802

**Authors:** Gregory J. Hunt, Mark A. Dane, James E. Korkola, Laura M. Heiser, Johann A. Gagnon-Bartsch

**Affiliations:** Department of Mathematics, William & Mary, USA; Department of Biomedical Engineering, Knight Cancer Institute, OHSU Center for Spatial Systems Biomedicine, Oregon Health and Science University, USA; Department of Statistics, University of Michigan, USA

## Abstract

The immediate physical and bio-chemical surroundings of a cell, the cellular microenvironment, is an important component of many fundamental cell and tissue level processes and is implicated in many diseases and dysfunctions. Thus understanding the interaction of cells with their microenvironment can further both basic research and aid the discovery of therapeutic agents. To study perturbations of cellular microenvironments a novel image-based cell-profiling technology called the microenvironment microarray (MEMA) has been recently employed. In this paper we explore the effect of preprocessing transformations for MEMA data on the discovery of biological and technical latent effects. We find that Gaussianizing the data and carefully removing outliers can enhance discovery of important biological effects. In particular, these transformations help reveal a relationship between cell morphological features and the extra-cellular-matrix protein THBS1 in MCF10A breast tissue. More broadly, MEMAs are part of a recent and wide-spread adoption of image-based cell-profiling technologies in the quantification of phenotypic differences among cell populations (Caicedo *et al*., 2017). Thus we anticipate that the advantages of the proposed preprocessing transformations will likely also be realized in the analysis of data from other highly-multiplexed technologies like Cyclic Immunofluorescence. All code and supplementary analysis for this paper is available at gjhunt.github.io/rr.

## 1. Introduction

The microenvironment of a cell encompasses its immediate physical and bio-chemical surroundings. This includes, for example, the adjacent extra cellular matrix (ECM), surrounding cells, ligands like hormones, cytokines, chemokines, growth factors, etc (Bhat and Bissell, 2014). These microenvironmental components modify cellular behavior through a host of different mechanisms. Accordingly, the interaction of cells with their microenvironment is a component of many cell and tissue level processes (Maman and Witz, 2018; Januschke and Näthke, 2014; Lin *et al*., 2012). For example, the extra-cellular matrix has been long known to regulate cellular functions like adhesion, migration, proliferation and differentiation (Teti, 1992; Bissell and Labarge, 2005; LaBarge *et al*., 2007). The microenvironment is also implicated in the development, progression, and ultimately treatment of many diseases and dysfunctions (Pelissier *et al*., 2014; LaBarge, 2013). For example, it has been posited that communication between B-cells and proximal stromal cells can promote malignant B-cell growth and drug resistance (Burger *et al*., 2009). Similarly, towards the goal of understanding therapeutic efficacy, it has recently been shown that the microenvironment of HER2-positive breast cancer cells can modulate drug response (Watson *et al*., 2018). Thus a better understanding of the microenvironment benefits not only basic research but also furthers an understanding of the interaction between therapeutic agents and regulatory behavior.

Understanding cellular microenvironments is part of the 2014 NIH Common Fund program called the Library of Integrated Network-Based Cellular Signatures (LINCS). The LINCS program is generally focused on understanding changes in cellular processes as a result of perturbing agents (Keenan *et al*., 2018). The Microenvironmental Perturbagen (MEP) program, one of six major branches of the LINCS program, specifically focuses on studying the influence of microenvironmental signals on cellular phenotype. To study signals in the microenvironment, researchers are using a novel high-throughput technology called the Microenvironment Microarray (MEMA)(Labarge *et al*., 2014; Watson *et al*., 2018; Lin *et al*., 2017; Smith *et al*., 2019). The technology, first developed by Mark LaBarge at Lawrence Berkeley National Laboratory (Lin *et al*., 2012; LaBarge *et al*., 2009), allows the study of several thousand combinations of microenvironmental factors on molecular and biological endpoints like cell proliferation, differentiation, or apoptosis. Specifically, MEMAs facilitate the study of these endpoints via high-throughput image-based profiling of cells.

A MEMA consists of a plastic substrate divided into several partitioned wells. Each well contains an array of several hundred ~ 400*μm* spots robotically printed onto the surface of each well. Thousands of cells are added to each well where they randomly bind to the spots. The cells on each spot interact with a pair of microenvironmental perturbagens. This perturbagen pair consists of an insoluble extra-cellular matrix protein (ECMp) and a soluble ligand. The ECMps are printed onto specific spots in the wells, while ligands are added to the buffer solutions in wells. Thus the cells in a spot interact with an ECMp specific to their spot and a ligand common to their well. In total, there are several thousand different combinations of ECMps and ligands perturbing the cells on a MEMA. Typically each perturbagen pair is replicated about a dozen times within each well. The cells are grown for several days and subsequently immunofluorescently stained and imaged with fluorescent microscopy. These images are analyzed with image-analysis software to produce quantified cellular features. These numeric quantifications of the cells’ properties constitute the data produced by a MEMA. The goal of a MEMA experiment is to analyze this data to further a biological understanding of the relationship between the perturbagen pairs and the physical properties of the cells (as quantified by the software-extracted features).

The features produced by MEMAs cover a wide range of cellular aspects. There are morphological features like cell area, compactness, eccentricity, perimeter, or solidity. The MEMAs also produce stain intensity features and features capturing the cell cycle state, cell lineage, cell count, texture and many others. Typically, several hundred features are extracted from the images taken of each spot.

The image features that are extracted from a MEMA can be flexibly adapted to suit the research interests. One can add to the list of extracted features by asking the software for more features or by using a different or more sophisticated image analysis program. Furthermore, since the underlying microscopy images are retained, features can be added retrospectively through re-analysis of the images with newer and better algorithms. Finally, the set of immunofluorescent stains applied to the cells dictates which features may be extracted. In turn, the stains applied depend on what physical structures are of interest. In summary, the number of features produced by a MEMA is not only large but the precise set of features will likely vary from one experiment to the next. This presents a problem for automatic processing of MEMA data. While the plethora cell-feature data presents new opportunities for discovery, it also necessitates an adaptive approach that can handle an ever-changing landscape of features.

In this paper we are interested in how best to transform the MEMA data to benefit downstream analysis. Data transformations are commonly part of the analysis of high-throughput -omics experiments. For example, microarray data typically is logarithmically transformed, RNA-seq data often is transformed as the log of one plus the read count, and mass cytometry data typically is transformed using a arc-hyperbolic-sine transformation. Unfortunately, determining a good transformation for MEMA data is more complicated than for many other classic -omics experiments due to the number, flexibility, and disparate nature of the features.

Importantly, there will not be a single transformation that works well for all features. For example, the appropriate transformation for the nuclei orientation will likely not be the same as that for the DAPI intensity. This follows because the two features have much different measurement scales and distributions. Orientation is approximately normally distributed with a mean of zero and intensity has a very right-skewed exponential-like distribution and is always positive (see Supplementary Figures 1 and 2). Thus we might, for example, want to transform intensity using a logarithmic transformation while we might not want to transform orientation at all.

The general problem is that the features produced by a MEMA each have their own different scales, distributions, ranges, outliers and quirks. There will not be one common transformation to apply to all the features. Rather, if there are a hundred MEMA features then we might potentially need a hundred different transformations. Unfortunately, it is not feasible to determine an appropriate transformation for each feature by hand, especially given that the set of features may vary from one experiment to another.

To overcome these problems, this paper comprehensively studies simple and robust ways of adaptively transforming MEMA features. We study ways of automatically transforming MEMA features to improve discovery of important biological effects and identification of unwanted latent technical effects. We demonstrate the utility of these transformations in both qualitative analyses (like data visualization) and quantitative analyses (like principal components analysis). In addition to enhancing analysis of features individually we show that these transformations are also beneficial when integrating features together. Towards this latter goal, we explore the interaction of data integration with data transformation. Finally, as part of all these analyses we employ methods to estimate principal components from data with missing values.

## 2. Data, Methods, and Motivation

In this section we briefly describe the structure of the MEMA data with which we work, outline our steps for processing the data, and motivate why these steps enhance visualization, integration, and discovery of latent biological and technical effects.

### 2.1. Structure of MEMA Data

In this paper we work with microenvironment microarray data from the MEP-LINCS Center at the Oregon Health and Science University. The data is accessible through Synapse with identifiers syn10155286, syn10155289 and syn10155292 (Gray *et al*., 2014). In total we analyze 24 MEMAs of human epithelial mammary tissue (MCF10A). The 24 MEMAs come in three batches of eight plates. Each MEMA plate is divided evenly into eight wells. Each well contains 700 spots in a 20 by 35 grid. Cells are added to the wells and bind to the spots. Subsequently, a buffer solution containing a specific ligand is added to each well. Thus the cells can grow out in the presence of different ECM proteins and ligands. The pattern of ECMps is identical across all wells (see Supplementary Figure 4) however a (potentially) different ligand is added to each well (see Supplementary Figure 5). After incubating the cells for 72 hours they are fluorescently stained, imaged, and cell-level features are extracted with image analysis software. For the analysis in this paper, we work with spot-level features (median summarized cell-level features). For each image feature we have a data matrix of 192 wells (3 batches × 8 plates × 8 wells) by 694 spots (we remove 6 alignment spots with no cells from the 700). In total we will work with 103 image features and thus have 103 feature matrices to process and analyze. See Supplementary Table 1 for a complete list of all features.

### 2.2. Robust Re-scaling

To process these feature matrices we follow three sequential transformation steps:

#### Procedure 1 Three-step Robust Re-scaling (RR)

**Step 1:** (G) robustly “Gaussianize” the data,

**Step 2:** (Z) convert the data to robust *z*-scores,

**Step 3:** (O) remove outliers.

These three steps are applied to each of the 103 feature matrices individually. In the following sections we explore the details of these steps and motivate why they enhance analysis.

#### 2.2.1. The Gaussianizing Step (G)

The (G) step transforms the data using a Box-Cox-like procedure (Box and Cox, 1964). We call this a “Gaussianizing” transformation because it makes the data approximately normally distributed. It first estimates a Gaussianizing (Box-Cox) transformation for each column of the feature matrix. It does this by optimizing over parameterized power and arc-hyperbolic-sine transformations and chooses the parameter for each column of the feature matrix that makes the column as normal as possible. Since it’s easy to over-fit these parameters, the procedure robustly chooses the median transformation parameter (across columns). We then apply the associated transformation element-wise to the feature matrix.

Let *Y* ∈ ℝ^*m×n*^ be a specific feature matrix. In our application we have 103 such feature matrices each with *M* = 192 rows (wells) and *N* = 694 columns (spots). Let 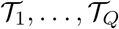 be a collection of *Q* parameterized transformation families so that for any *q* = 1,…,*Q* the family 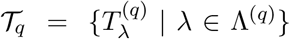 consists of differentiable, monotonic, transformations 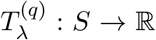 on some *S* ⊆ ℝ. The goal is to optimize over the union of these families and choose the transformation that makes the data close to being normal (without over-fitting).

In this manuscript we choose *Q* = 2 families over which to search: a power family

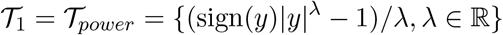

and an arc-hyperbolic-sine family

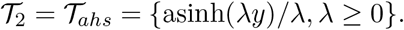

We choose these two families because they cover a range of power and sigmoidal shapes. Many reasonable choices of parameterized families can be made and nothing in our discussion depends on the specific choices. We include other options of families in our software.

Before we describe the procedure for optimizing over many families, we will first consider the simpler case when *Q* = 1 and discuss how to choose an optimal transformation over a single family generically denoted 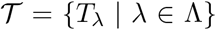. Define *Y*_\**j*_ be the *j^th^* column of *Y*, and for any λ ∈ Λ let *Y*_\**j*_ (λ) = *T*_λ_(*Y*_\**j*_) be *Y*_\**j*_ under the transform *T*_λ_. The goal is to choose a 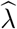 so that 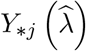 is approximately normally distributed for each *j* = 1,…,*N*. The (G) approach follows two steps:

For the first step, estimate 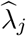 using the traditional Box-Cox approach on *Y*_\**j*_. Assume there is some λ_*j*_ so that 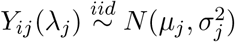 for *μ_j_* ∈ ℝ and *σ_j_* ≥ 0. Then let 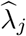 be the MLE of λ_*j*_. This is obtained by profiling the likelihood over *μ_j_* and 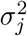 and then maximizing the profile likelihood over λ_*j*_. If *L_j_* is the profile likelihood of λ_*j*_ profiling over *μ_j_* and 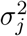 then

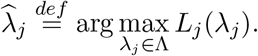

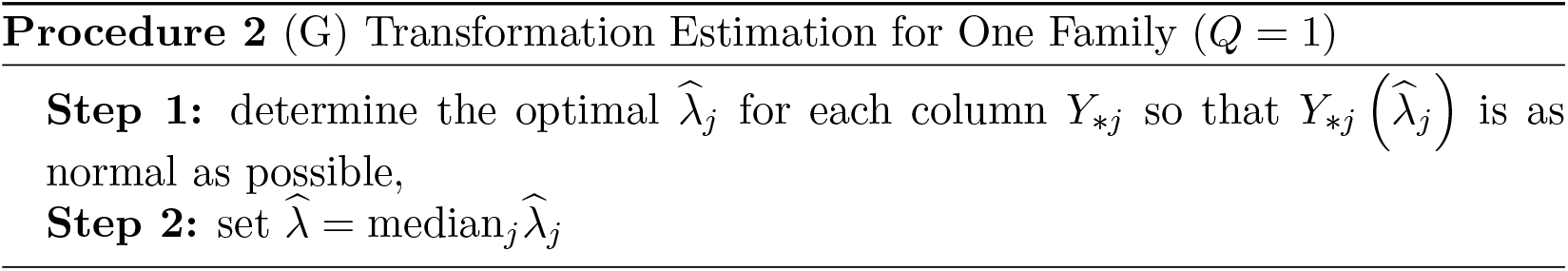

After estimating 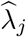 for each column, the second step is to summarize the collection of 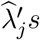 into a single 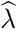. This is done with the median. Define 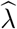 as the element-wise median 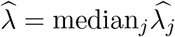.

To summarize, the procedure when *Q* = 1 is to first optimize within each column and then median-summarize across the column-wise estimates. When *Q* > 1 we add an additional step to first determine which family among 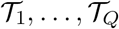 is best. The procedure is described in Procedure 3.

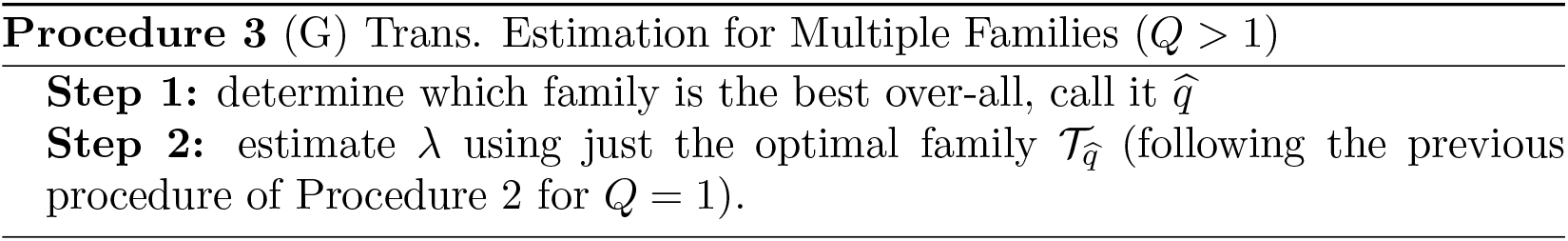

This procedure first determines the best family individually for each column and then uses the family that is best among a plurality of columns. More specifically, let *L_j_*(*q_j_*, λ_*j*_) be the likelihood of the *j^th^* column after transformation using the 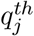 family and transformation parameter λ_*j*_. Optimize *L_j_*(*q_j_*, λ_*j*_) jointly over λ_*j*_ and *q_j_* and let

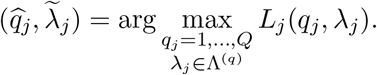

and

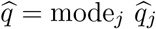

so that 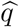 is the family that is the best among a plurality of the columns. Once we have determined this optimal family 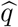 we then estimate 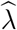 following the procedure when *Q* = 1 using the family 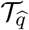. Finally, define the Gaussianized version of *Y* as

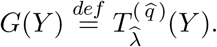

#### 2.2.2. The z-score Step (Z)

The second step in the (RR) procedure is the (Z) step, a robust *z*-score transformation. Let 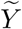 be a vectorized version of *Y* and 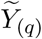 be the *q*-winsorized version of *Y*. Jn this paper we will use *q* = 0.001 replacing everything below the *q^th^* quantile of 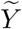 by the *q^th^* quantile and replacing everything above the (1 − *q*)^*th*^ quantile of 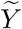 by the (1 − *q*)^*th*^ quantile. Given this, the robust *z*-score version of *Y* is defined as

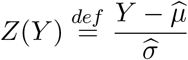

where 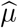 and 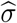 are mean and s.d. estimates of 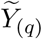.

#### 2.2.3. The Outlier Removal Step (O)

The final of the three (RR) steps is outlier removal. The outlier removal procedure simply thresholds *z*-scores and marks as missing anything beyond four standard deviations. First let *Z*(*Y*) be the robust *z*-scored version of *Y*. We then define *O*(*Y*) as

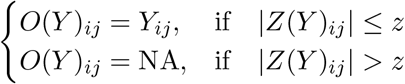

where “NA” denotes a missing value. To be conservative in this paper we use *z* = 4 although this is somewhat arbitrary. With *z* = 4 if the data is truly normal this removes only about 3e-3 percent of the data from each tail.

#### 2.2.4. The Three-Step (RR) Procedure

Given these definitions, the three step (RR) transformation is to apply the (G), (Z), and then (O) transformations. If *Y* is a feature matrix then we define *RR*(*Y*) as

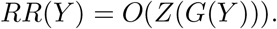

In the remainder of this section we will motivate (1) why these steps help discovery of important latent effects in the data and (2) how this processing improves data integration.

### 2.3. Motivation: Discovery of Important Latent Effects

A central component in the analysis of MEMA data is the identification of important latent effects. We divide these latent effects into two main categories: (1) biological effects and (2) technical effects. Biological effects are the effects of primary research interest. Such effects include, for example, differences in biological endpoints due to ECMps or ligands. On the other hand, technical effects are unwanted and we are interested in identifying them so that we may remove them. Examples of these effects include batch across plates or wells and spatial effects within wells. Discovery of either technical or biological latent effects is typically done through visual inspection of plots or quantitative analysis like PCA. Unfortunately, methods like PCA are often misled by prominent aspects of the data that tell us little about either technical or biological effects.

As an example, consider how PCA can be misled when used to identify groups in skewed data. Let *u*^(1)^, *u*^(2)^, *v*^(1)^, *v*^(2)^ ∈ ℝ^*N*^ have elements that are *i.i.d* from a standard log-normal distribution. For a small *δ* ∈ ℝ and noise *ϵ* ∈ ℝ^2*N*×*N*^ define a block data matrix *Y* as

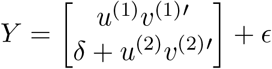

so that the first *N* rows of the data matrix and the last *N* rows of the data matrix constitute two groups with a mean difference of *δ*. The left side of Figure 1 displays a histogram of the elements of *Y* for a simulation using *δ* = 1/2 and *ϵ* distributed *i.i.d*. standard normal. It is difficult to distinguish between the two groups in the right-hand panel of Figure 1 because the group difference is over-shadowed by the data’s long tails. Consequently, PCA identifies the variance due to tail skewness, not the group difference, as the most prominent variation in the data. While the first two principal components capture more than 99% of the total variance in this example, they only capture about 50% of the group difference (see Supplementary Figure 3). The right side of Figure 1 shows that a log transformation makes the groups more prominent. The transformation un-skews the data thereby attenuating the effect of the tails on PCA. In this case, while the first two PCs only capture about 80% of the total variation, they capture about 94% of the variation due to group difference (see Supplementary Figure 3).

**Fig. 1.**
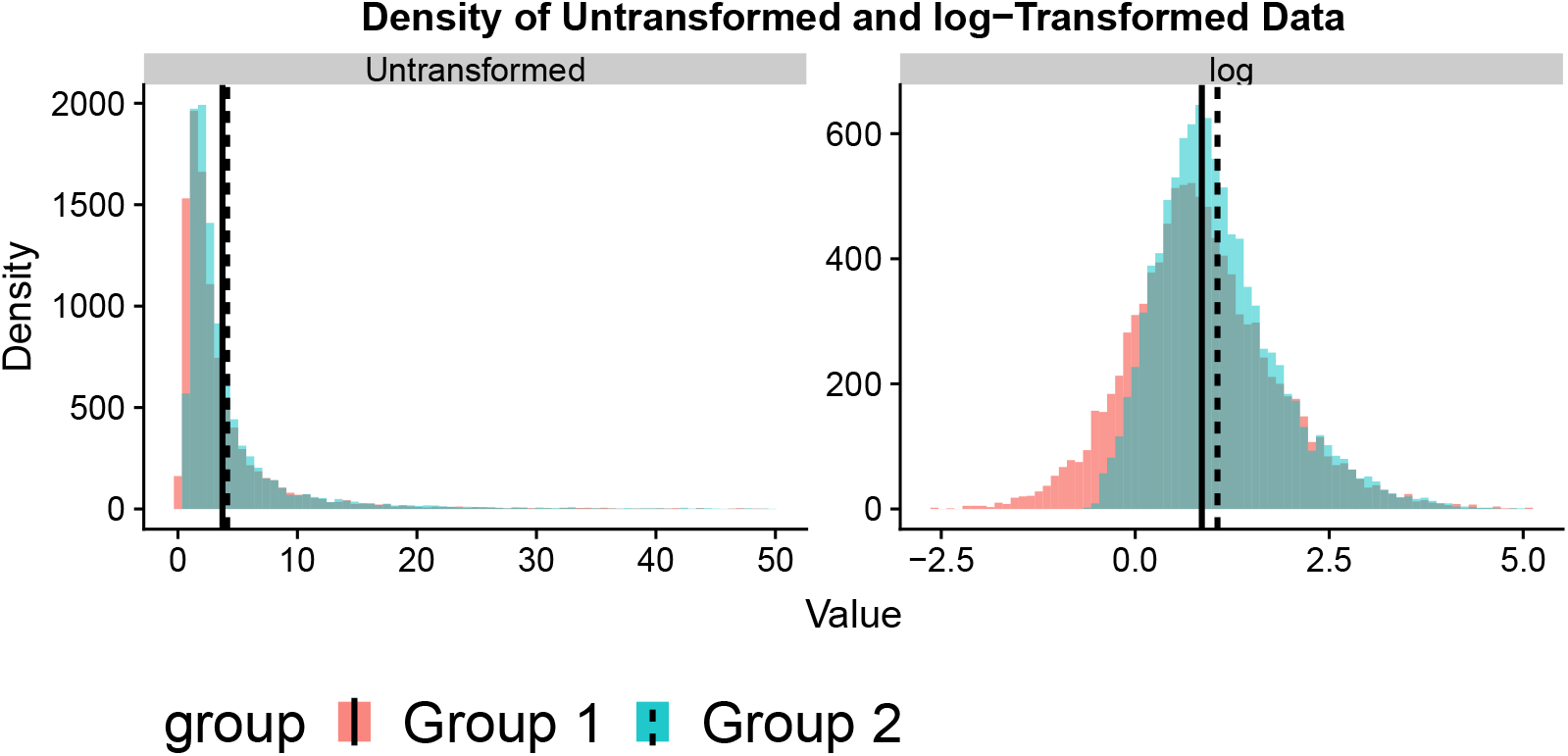
(A) The percentage of cumulative variance captured by first *k* principal components for both un-transformed data and log-transformed data. (B) The mean squared canonical correlations between the grouping factor and the first *k* principal components.

As motivated by the previous example, we want to attenuate the influence of prominent, yet uninformative, variation. Our processing steps described in Section 2.2 attempt to ameliorate the effects of two commonly encountered, and potentially misleading, aspects of MEMA data. Those aspects are (1) skewness in measurement scales and (2) anomalous outliers. By anomalous outliers we mean extremely unusual data points that are not informative of much beyond their own uniqueness. These outliers could result from biological or technical artifacts. For example, cells may have difficulty growing on a spot, distorting the median-summarized morphological measurements of the cells present. Or, for example, if the image-analysis software has difficulty determining the boundary between overlapping cells leading to features that do not properly reflect the true biology.

To guard our analysis against un-interesting variation we follow the three robust re-scaling steps (G), (Z) and (O) outlined previously. The (G) step is used to prevent a feature’s naturally long-tailed measurement scale from dominating analysis. This is done by applying a robust Box-Cox-like procedure to “Gaussianize” the data and adaptively de-skew each feature’s distribution. We use (G) instead of the traditional Box-Cox procedure as the latter will be highly influenced by outliers. Box-Cox will propose an extreme transformation to rectify single outliers that it perceives as a huge violation of normality. Conversely, (G) is “robust” to outliers as it estimates 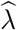 as the median of the column-wise 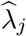 estimates. (G) only transforms the data to un-skew fundamentally skewed data not simply to reign in a few points. Dealing with outliers is more parsimoniously done by specific outlier targeting procedures not a global Gaussianizing transformation. To remove outliers (RR) first converts the data to robust *z*-scores using (Z) and then removes any entry of the feature matrix bigger in magnitude than four (the (O) step).

### 2.4. Complete Singular Vectors

The primary tool we use to recover latent effects is the singular value decomposition (SVD). For a feature matrix *Y* with a singular value decomposition (SVD) of *Y* = *UΣV*′ we call the columns of the *U* left singular vectors and the columns of *V* the right singular vectors. When the columns of *Y* have been mean-centered the left singular vectors are often called the principal components (PCs). We avoid this terminology because we do not mean-center.

When calculating the SVD for MEMA data we need to account for missing data. Missing values arise for biological reasons (e.g. the cells failed to grow), image-analysis reasons (e.g. the software could not detect any cells), and because (RR) introduces missing values as part of (O). As the SVD is undefined for matrices with missing values, we use “complete” singular vectors calculated from re-scaled pairwise-complete gram matrices. Note that this is similar to the pairwise-complete option for `cor` in `R` (R Core Team, 2018).

Let *Y* ∈ ℝ^*M×N*^ be a feature matrix with missing values. Define *Y*_0_ as *Y* with missing values replaced by zeros and 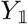 so that 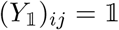 {*Y_ij_* is not missing}. We define the re-scaled pairwise-complete left Gram matrix *Y · Y*′ so that

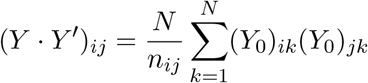

where 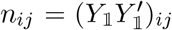 is the number of pairwise non-missing entries between row *i* and *j* of *Y*. This is well-defined so long as none of the entries of 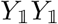 are zero.

The matrix *Y · Y*′ is a matrix of re-scaled inner products of the rows of *Y* accounting for the number of non-missing pairs between rows. We similarly define the right gram matrix *Y′ · Y* replacing *Y* for *Y*′ above. We call the eigenvectors of *Y · Y*′ the complete left singular vectors and the eigenvectors of *Y′ · Y* the complete right singular vectors. We order these vectors decreasing by associated eigenvalue keeping only those associated with positive eigenvalues. For brevity we often omit the adjective “complete”, referring to these simply as the “singular vectors.” If there are no missing values they are one and the same.

### 2.5. Average Singular Vectors

In addition to recovering important latent effects in individual features, we are interested in latent effects common to multiple features. To extract a common set of latent effects from a collection of *P* features *Y*^(1)^,…, *Y*^(*P*)^ we use the eigenvectors from the average of their re-scaled pairwise-complete left and right gram matrices, respectively,

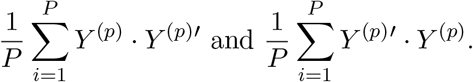

We call these eigenvectors the left and right average singular vectors (ASVs), again retaining only those eigenvectors associated with positive eigenvalues.

## 3. Results

### 3.1. Features and Transformations Considered

The MEMA plates we analyze are grown, stained and imaged in three separate processing batches. A different set of stains is used in each batch. Those sets are: (1) “SS1” (containing stains DAPI, Actin, CellMask and MitoTracker), (2) “SS2noH3” (containing stains DAPI, Fibrillarin and EdU), and (3) “SS3” (containing stains DAPI, KRT5, KRT19 and CellMask). Because each of these batches use a different staining set we refer to them as the “staining batches.” While these batches are separate experiments, aside from the staining set the experimental conditions were made as identical as possible.

In total there are 103 different image features extracted from the MEMAs. A different set of features is extracted in each staining batch with some being common across multiple batches. There are 50 features extracted in at least two of the staining batches and 18 features that are extracted from all three. We focus on four features in this paper:

a. cell area (notated on synapse as “Cells_CP_AreaShape_Area”)
b. cell compactness (“Cells_CP_AreaShape_Compactness”)
c. spot cell count (“Spot_PA_SpotCellCount”)
d. total cytoplasm DAPI intensity (“Cytoplasm_CP.Intensity _IntegratedIntensity_Dapi”).

We choose these features because they represent several different feature types. The first two are morphological traits of cells, the third is the cell count, and the last is an intensity. We have deposited the full results for all features at gjhunt.github.io/rr.

To explore the effects of (G), (Z) and (O), we consider five transformations of the features: (1) no transformation (NT), (2) the (G) step only (3) the (Z) step only (4) the (O) step only (5) the three-step (RR) transformation.

### 3.2. Visualization

#### 3.2.1. Feature Distributions

A typical first step in exploratory analysis is data visualization. Simple data visualizations can succinctly summarize the major features of the data and inform qualitative analyses. In Figure 2 we plot the distribution of cell area for the five transformations. The colored densities correspond to staining batches. The black line is the density of all data combined. Notice in Figure 2 that the density of (NT) largely reflects the data’s long tail. The same can be said for (Z). Conversely, the other transformations reveal the staining batches. Both (O) and (G) de-emphasize the data’s long tail in favor of the group difference. Furthermore, in (RR) these groups are approximately Gaussian.

**Fig. 2.**
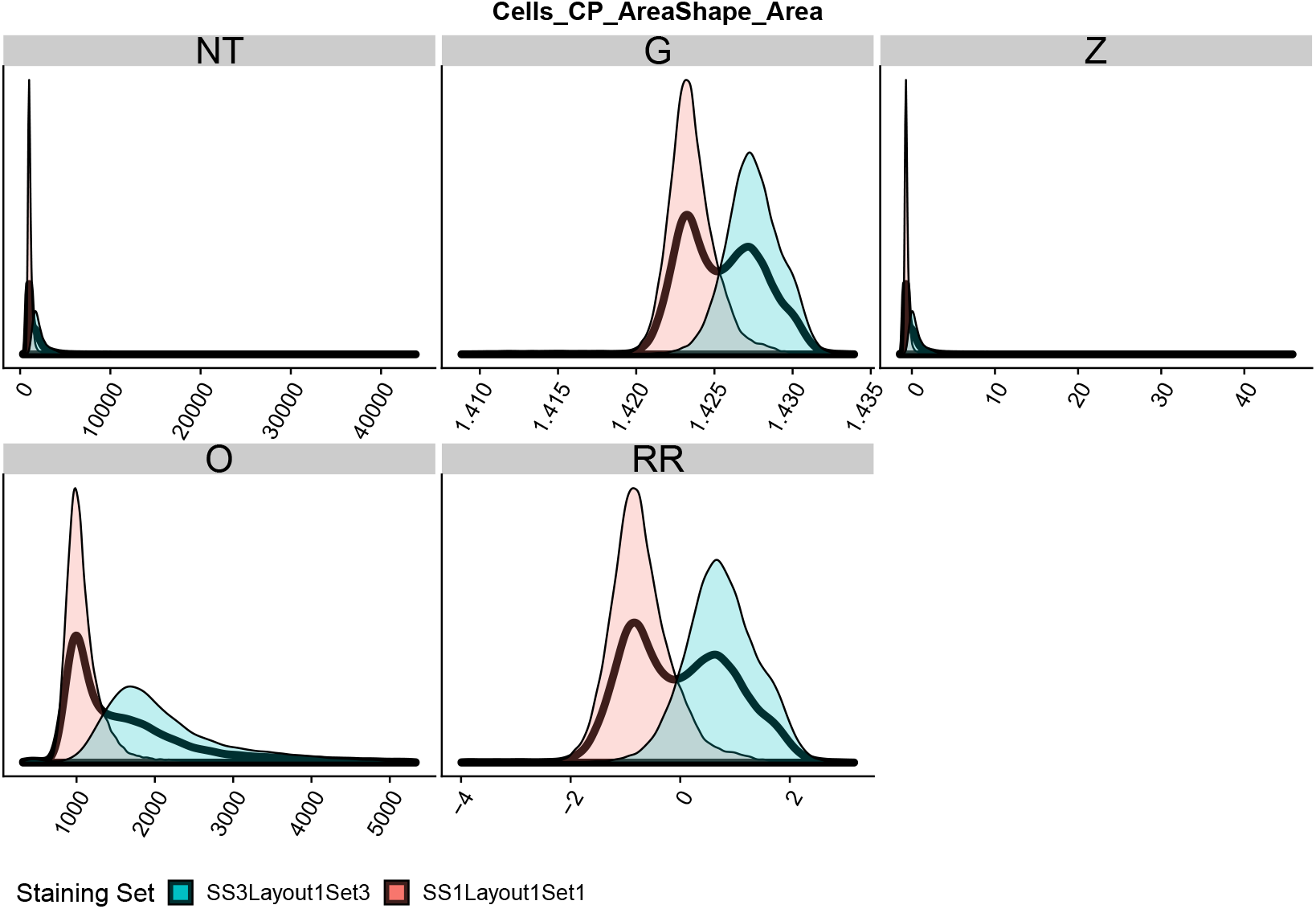
Density of elements of cell area feature matrix. Black density is all elements combined. Colored densities are the densities for the two staining batches. Subplots are for five processing transformations of this matrix: (NT) no transformation, (G) Gaussianization, (Z) *z*-score, (O) outlier removal, (RR) the three-step (G), (Z), and (O), robust re-scaling.

Supplementary Figure 6 shows a similar plot to Figure 2 but for the other example features. Largely we see the same behavior. In Supplementary Figures 7, 8 and 9 we display density plots similar to Supplementary Figure 6 but for different wells, plates, and ligands instead of the staining batches. These other covariates are picked up in a similar, albeit more attenuated, manner.

#### 3.2.2. Heat-maps

Another way to visualize the MEMA data is through heat-map pseudo-images. These pseudo-images are heat-maps of the value of a feature for each spot plotted following the same physical layout as the MEMA. These pseudo-images can be useful for discovering spatial effects and assessing the quality of data. As an example, we visualize cell area this way in Supplementary Figure 10. In Figure 3 we display a single well from Supplementary Figure 10 across the five transformations. The colors are more blue if they are close to the minimum cell area, red if they are close to the maximum, and white if they are half-way between. Dark grey spots are omitted according to the MEMA design. Note that the color scale is determined globally over all wells and all plates in the dataset (see Supplementary Figure 10).

**Fig. 3.**
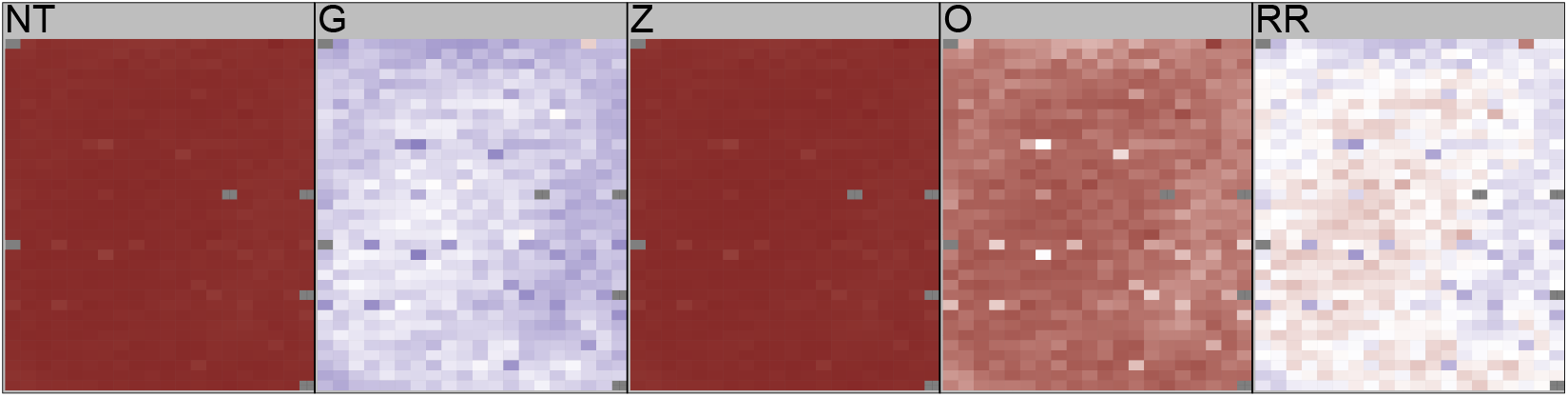
Heat map of a single well across the five transformations (NT), (G), (Z), (O), (RR). This is a sub-plot of Supplementary Figure 10. Color scaled is determined globally over all spots, wells, and plates in the dataset to reflect the fact that the transformation is similarly calculated over this data. Thus we see no blue in this (NT) sub-plot as we see almost no blue in Supplementary Figure 10. This plot is a representative microcosm of the larger plot.

This figure is not very informative for (NT) or (Z). The skewness and outliers assign the bulk of data to a tiny range of colors meaning the plots are essentially a single color. Conversely, for (G) and (O) we see a spatial effect between the right and edges and the rest of the well. We also see a non-spatial effect where certain spots are much different than their surroundings. We circle these spots in orange in Supplementary Figure 11. In Section 3.5 we show that this is an effect of the ECMps NID1 and ELN. We can see from these plots that (RR) strongly highlights the spatial effects as well as the NID/ELN effect.

In Figure 4 we focus on a different well of Supplementary Figure 10. Here, the green spots indicate missing data. These spots are missing either due to experimental error or because they have been removed as part of analysis. We see similar behavior where (G) and (O) reveal spatial differences between the upper right and the rest of the well. This spatial effect is also seen in (RR) however the number of points removed is much different in (RR) compared to (O). This highlights the difference between (O) thresholding outliers without transformation and (RR) thresholding outliers after (G). We believe thresholding based on a *z*-score makes most sense on a Gaussianized scale (as in (RR)). Note also that outliers are defined in a global context of the entire data, so while many values are marked as outliers by (RR) in this particular well it is a small percentage of the entire data.

**Fig. 4.**
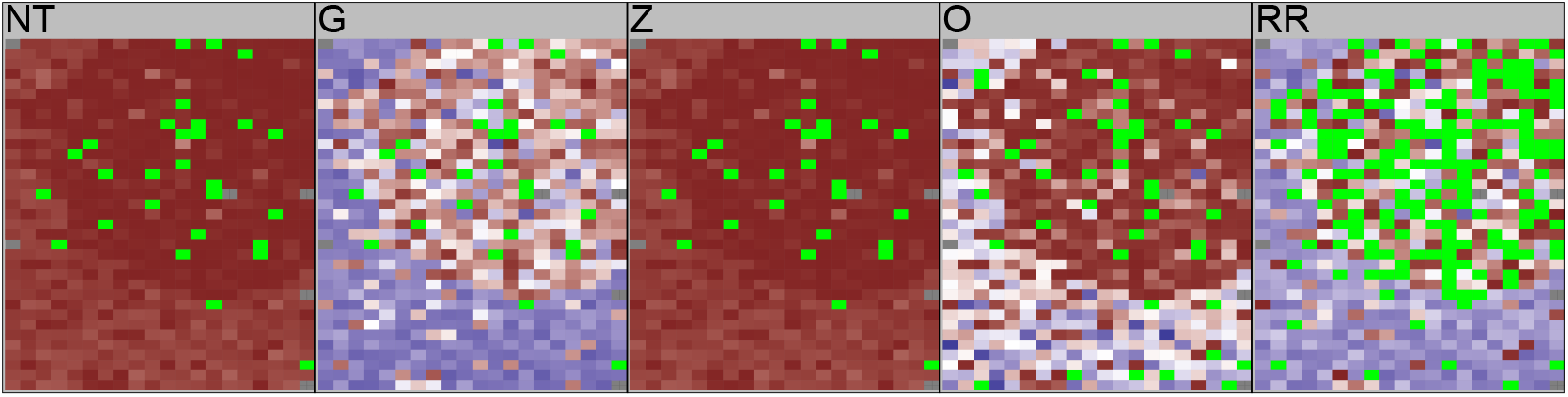
Similar to Figure 3 but focusing on a different well.

In addition to highlight spatial effects, these transformations also reveal batch effects between plates, wells, and staining batches. In Figure 5 we display the heat-map pseudo-image of cell area for eight wells across (NT), (G), (O) and (RR). ((Z) is identical to (NT).) The top four wells in each sub-plot are from the first staining batch, the bottom four wells are from the second. Nonetheless, we see little indication of batch in (NT). However batch is visible in (G), (O) and (RR). The bottom of (G) is lighter blue than the top, and the top of (O) is lighter red than the bottom. In (RR) we have solid-blue in the top and mostly red in the bottom. Being better able to identify batch effects hopefully will aid down-stream procedures to account for such effects.

**Fig. 5.**
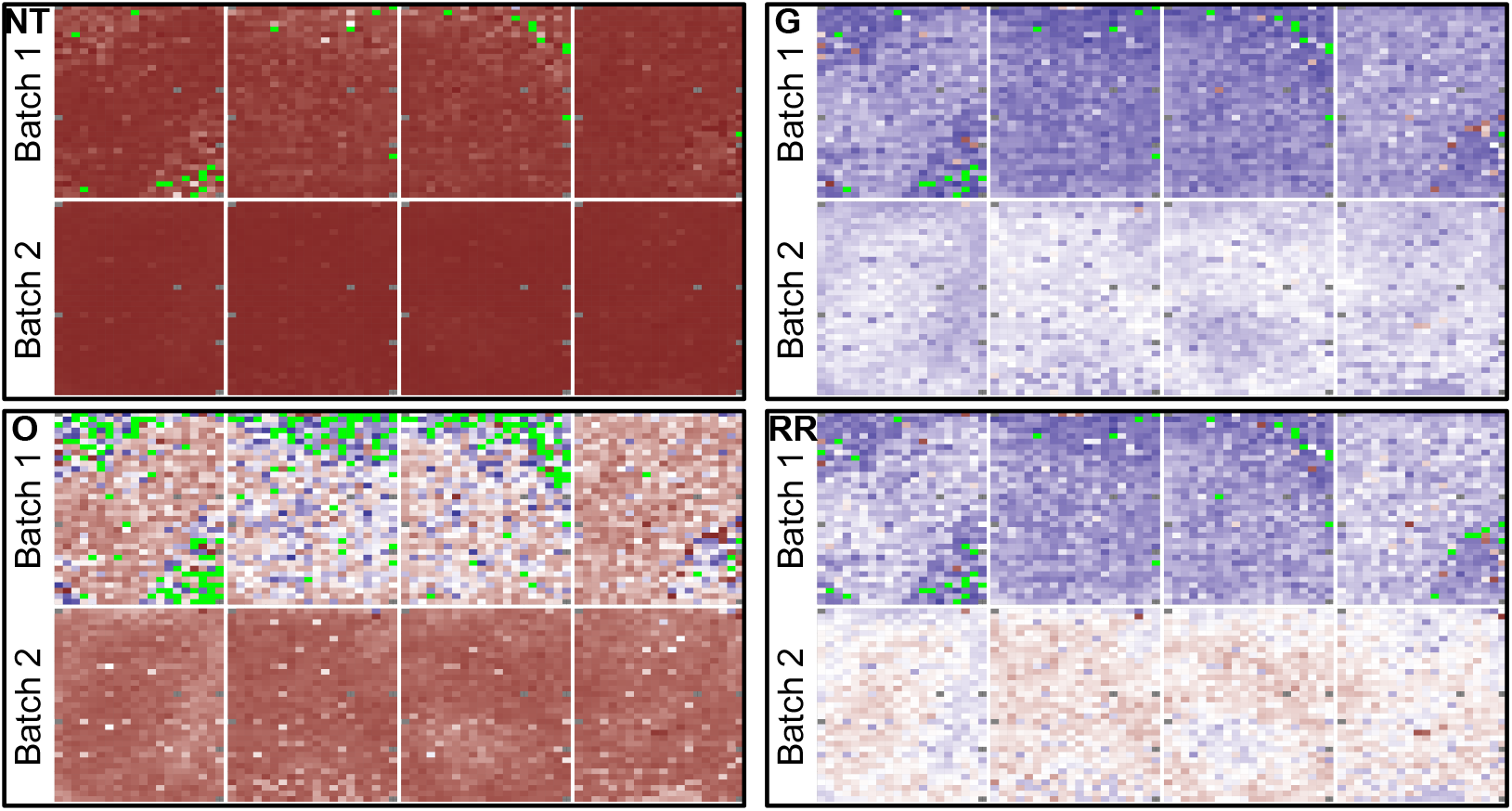
Heat map of a eight wells across the five transformations (NT), (G), (Z), (O), (RR). Top row of each subplot is from first staining batch. Bottom row is from second staining batch. Colors are more blue if they are close to the minimum, red if they are close to the maximum, and white if they are close to half-way between. Green spots are missing. Dark grey spots are omitted according to the MEMA design.

### 3.3. Recovering Technical Effects Across Wells

Batch effects are a common and well-studied problem in high-throughput biological experiments like MEMAs (Leek *et al*., 2010). Often, such batch effects obscure biological variation of interest. To deal with this problem, unwanted variation like batch is typically identified using the SVD and projected out of the data. In this section we explore how (RR) helps identify unwanted variation like batch using the SVD. We focus on the large staining batch effect as it was visible by eye in Figure 5.

We assess the transformations by measuring the percentage of the batch captured by the first *k* singular vectors of the transformed feature matrix. Let *U* = [*u*_1_,…, *u_N_*] ∈ ℝ^*M×N*^ be the (complete) left singular vectors of a feature matrix and *B* ∈ ℝ^*M×D*^ be the batch indicator matrix so that *B_ij_* = 1 if well *i* is in batch *j* for *j* = 1,…, *D*. Here we have *D* = 3 for the three staining batches. For *k* = 1,…, *N* and *t* = 1… min(*k, D*) define 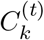 to be the *t^th^* canonical correlation between the first *k* left singular vectors *U_k_* = [*u*_1_,…,*u_k_*] and the batch *B*. Then let

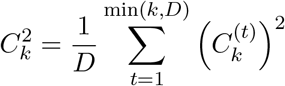

to be the average of these squared canonical correlations. We can interpret 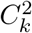 as the percentage of the batch *B* that is captured by these first *k* singular vectors. In Figure 6 we plot 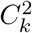 on the *y*-axis and vary *k* across the *x*-axis from *k* = 1 to 192.

**Fig. 6.**
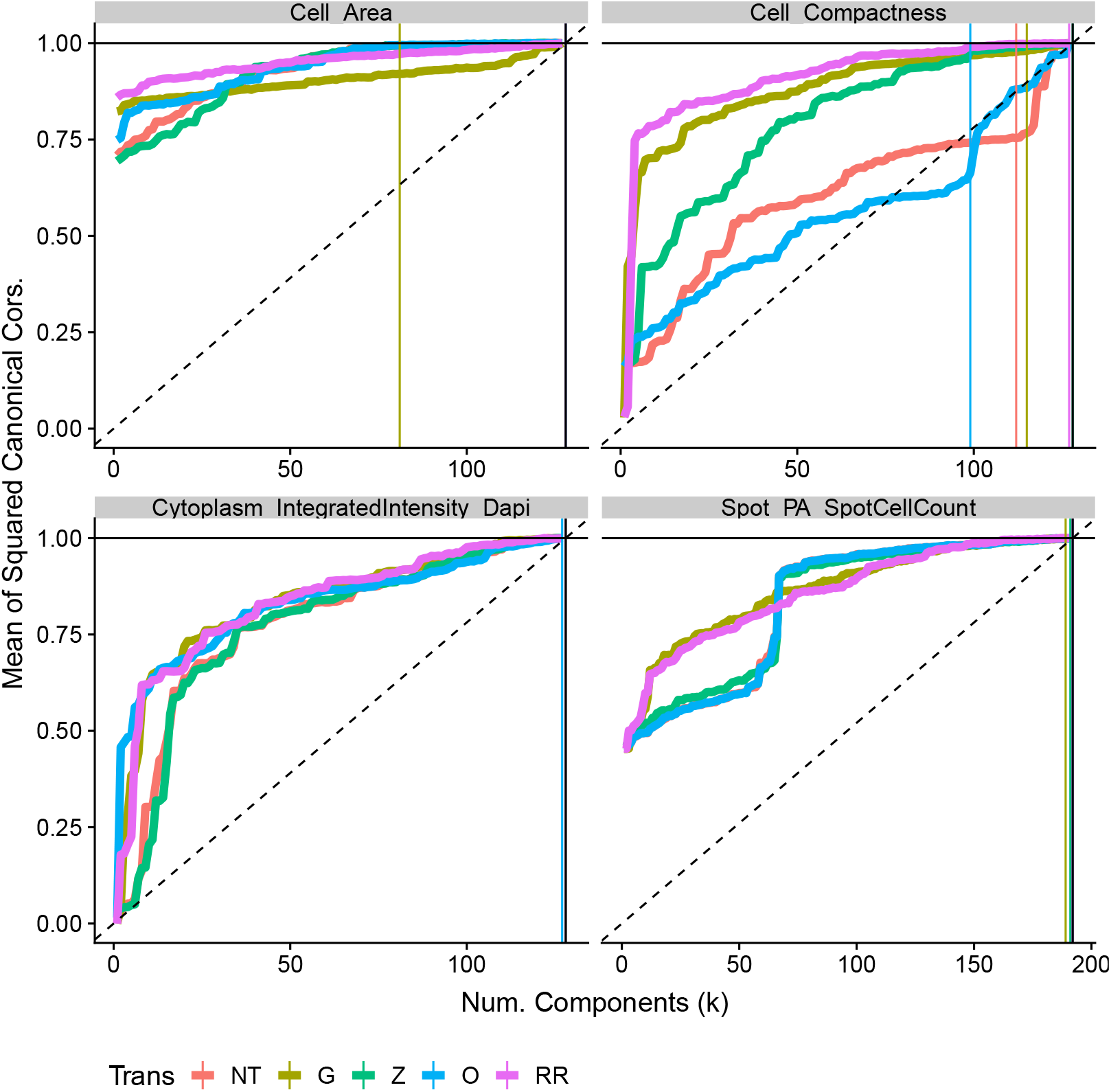
Mean of the squared canonical correlations between the first *k* left singular vectors and the staining batch dummy variables.

From this figure we see that the transformations enhance identification of the staining batch. Consider the cell area and total DAPI intensity features. As compared with no transformation (NT), these plots shows that (G), (O), and (RR) increase how much of the staining batch is captured by the first several singular vectors. These transformations attenuate the non-informative tails of the distributions and focus the singular vectors on the differences across the staining batches.

We summarize batch recovery for all features in Figure 7. Here, we calculate the area under the CC curves (AUC) for each feature as 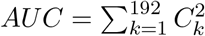 Broadly, we see the same behavior in Figure 7 as displayed in Figure 6. (RR) seems to generally improve recovery of the staining batch. Sometimes we see a substantial improvement (e.g. Cell Compactness) and rarely do we see that (RR) is detrimental.

**Fig. 7.**
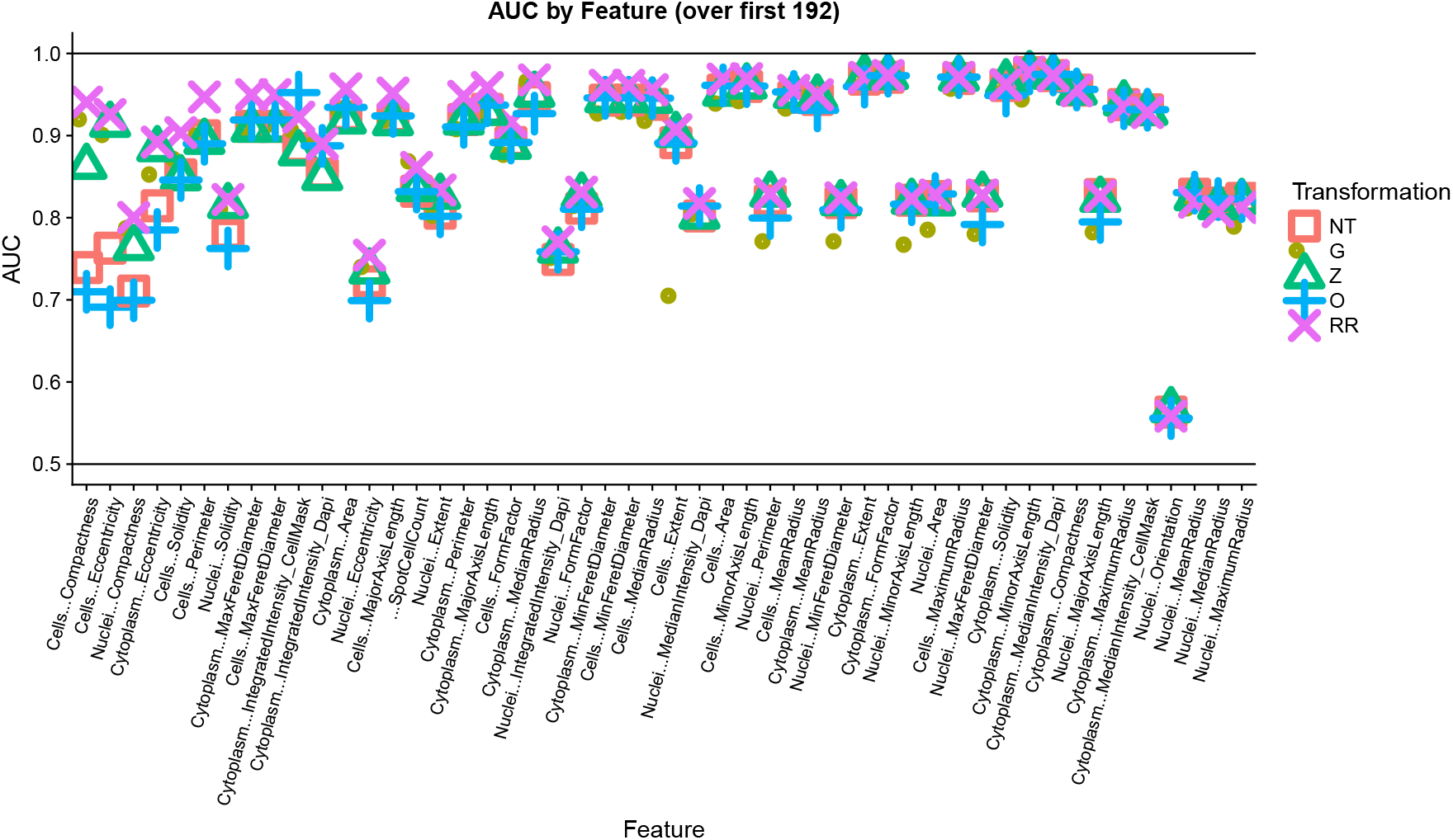
Grand mean of the squared canonical correlations across number of components (*k*). Canonical correlation is calculated between the first *k* left singular vectors and the staining batch dummy variables. Variable names have been shortened for readability. We order the features left to right decreasing by the difference in the AUC between (RR) and (NT). Thus those on the left are where (RR) performs relatively better than (NT) and less so on the right.

In Supplementary Figures 15 - 20 we display similar plots for the recovery of plate, well, and ligand effects. While these effects are not as prominent, we still see that the (RR) transformation slightly improves recovery of these latent effects without being detrimental.

### 3.4. Data Integration for Discovering Between-Well Effects

Given the close relationship among many of the MEMA image features, latent effects that appear in one feature may show up in other features. Shared effects give insight into biological and technical aspects that are important across many features. To extract these common effects we integrate information across MEMA features using the left average singular vectors (ASVs) as described in Section 2.

In the left panel of Figure 8 we plot the mean squared canonical correlations 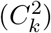 between the first *k* left ASVs and the staining batch. We calculate the ASVs using the 18 features measured in every plate. From this figure we can see that (Z) and (RR) quickly and strongly recover the staining batch. The AUC for these curves is in excess of 0.95, meaning it recovers batch better than the majority of individual features. The ASVs “average-out” feature-specific effects and amplify common effects like staining batch.

**Fig. 8.**
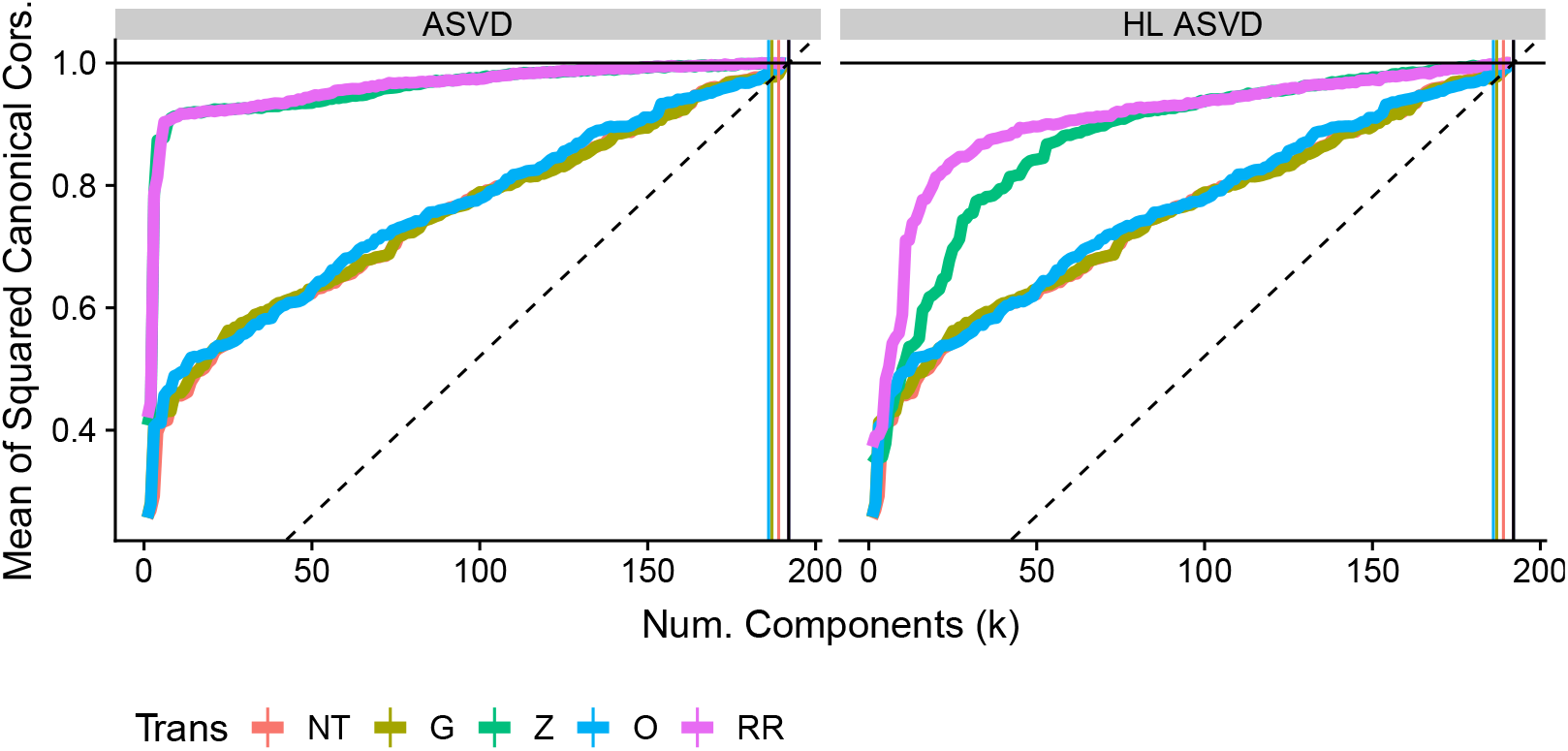
Mean of the squared canonical correlations between the first *k* average left singular vectors and the staining batch dummy variables. The average left singular vectors come from integration of (Left) the 18 features that are measured across all MEMAs, and, (Right) the five with the highest leverage points (among those 18).

It is notable that the (Z) and (RR) recover batch significantly better than (O), (G), and (NT). This happens because the ASVs element-wise average Gram matrices across features. If these Gram matrices are on vastly different scales their average is biased towards the largest features. This arbitrarily weights features’ by their scales. To equitably integrate information all features should have values in a similar range as in (Z) and (RR). Thus they recover batch better.

Finally, it is notable how well (Z) does alone. This happens because the averaging used to compute the ASVs conveys some of the same benefits as (G) and (O). This is true so long as we do not have a small number of features or systematic skewness or outliers across features. In the right panel of Figure 8 we calculate the ASVs using only five features with several high-leverage points. Here we see a separation between (Z) and (RR) since the average is over a small number of highly-skewed features. In any case, including (G) and (O) steps does not seem to hurt the analysis and thus we still recommend the full three-step (RR) transformation for integrating features in this manner. Similar, but attenuated results for plate, well and ligand are shown in Figures 21 - 23.

### 3.5. Discovering Biological and Spatial Effects within Wells

The left singular vectors of the feature matrices reveal latent effects across the wells, plates, and staining batches. Similarly, the right singular vectors (RSVs) reveal effects across the spots. In Figure 9 we display a scatter plot of the first four RSVs of total cytoplasmic DAPI intensity for (NT) and (RR).

**Fig. 9.**
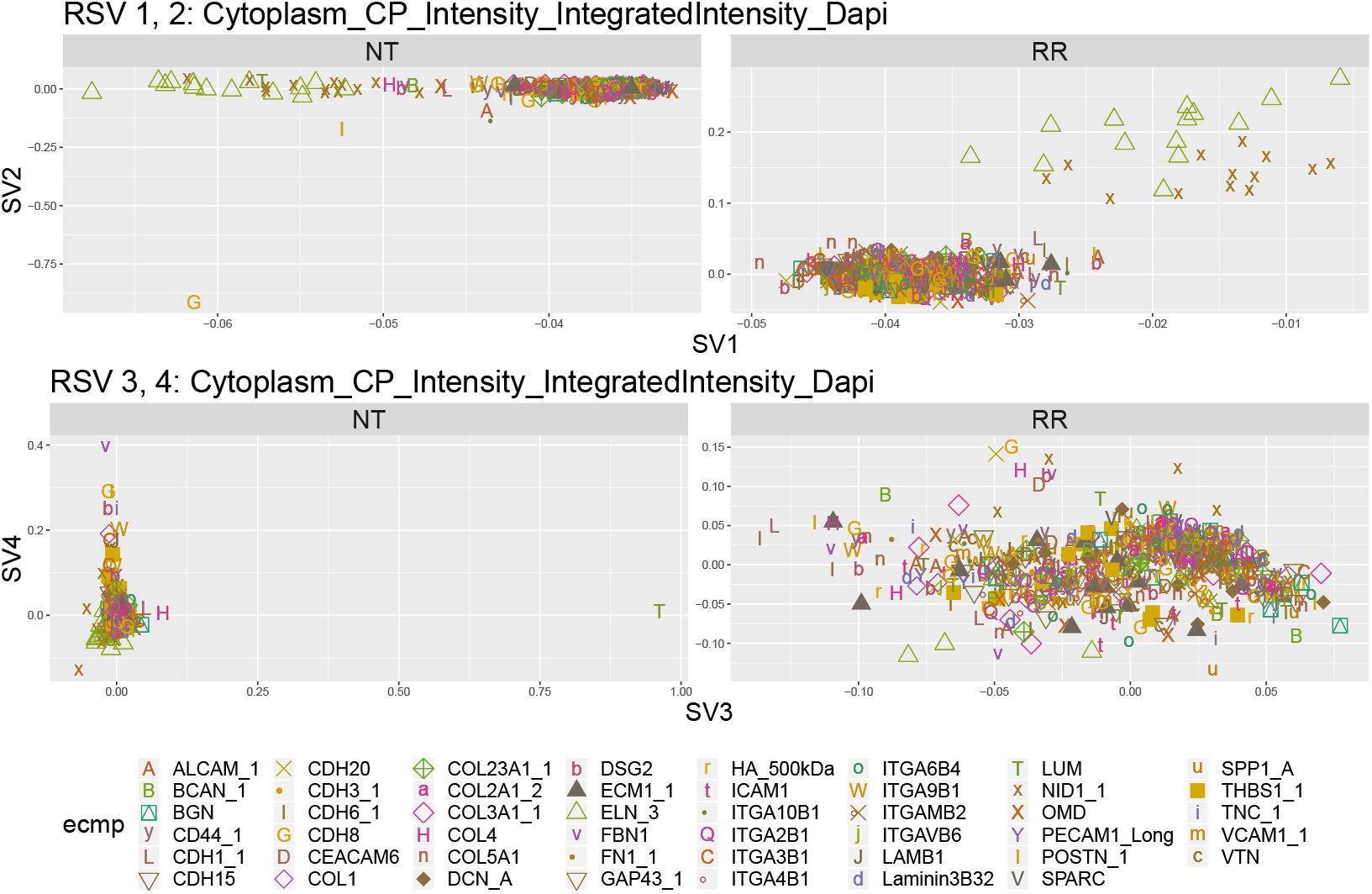
Scatter plot of elements of first four right singular vectors against each other for the total cytoplasmic DAPI intensity feature. Shape and color indicate ECMp of the spot corresponding to the elements of the singular vector.

A prominent feature of Figure 9 is the separation between the ECMps ELN, NID1, and the rest. Upon further investigation of the underlying MEMA images we find that this effect manifests because the cells have difficulty adhering to the ELN and NID1 ECMp substrates. Notice the cell count heat-map in Supplementary Figure 13 shows that the cell count in the ELN and NID1 spots are significantly lower than other spots. While this ELN-NID1 effect is present in the un-transformed data, it is more prominent in (RR). The first RSV from the un-transformed data does capture the effect, however the second through fourth RSVs are focused on explaining several outliers. Moreover, (RR) separates NID1 and ELN from the other ECMps and from each other.

In Figure 10 we plot pseudo-image heat-maps of the first ten RSVs for cytoplasmic DAPI intensity arranging elements of the RSVs according to the MEMA plate spatial layout. In addition to the ELN/NID effect, these plots reveal common spatial patterns across wells. These patterns are more visible for (RR) than (NT) as the RSVs of (NT) mostly capture outliers. It is important to identify such unwanted effects so that we can properly account for them downstream. In Figures 25 - 29 we display similar scatter plots and heat-maps for the other example features. They tell similar stories.

**Fig. 10.**
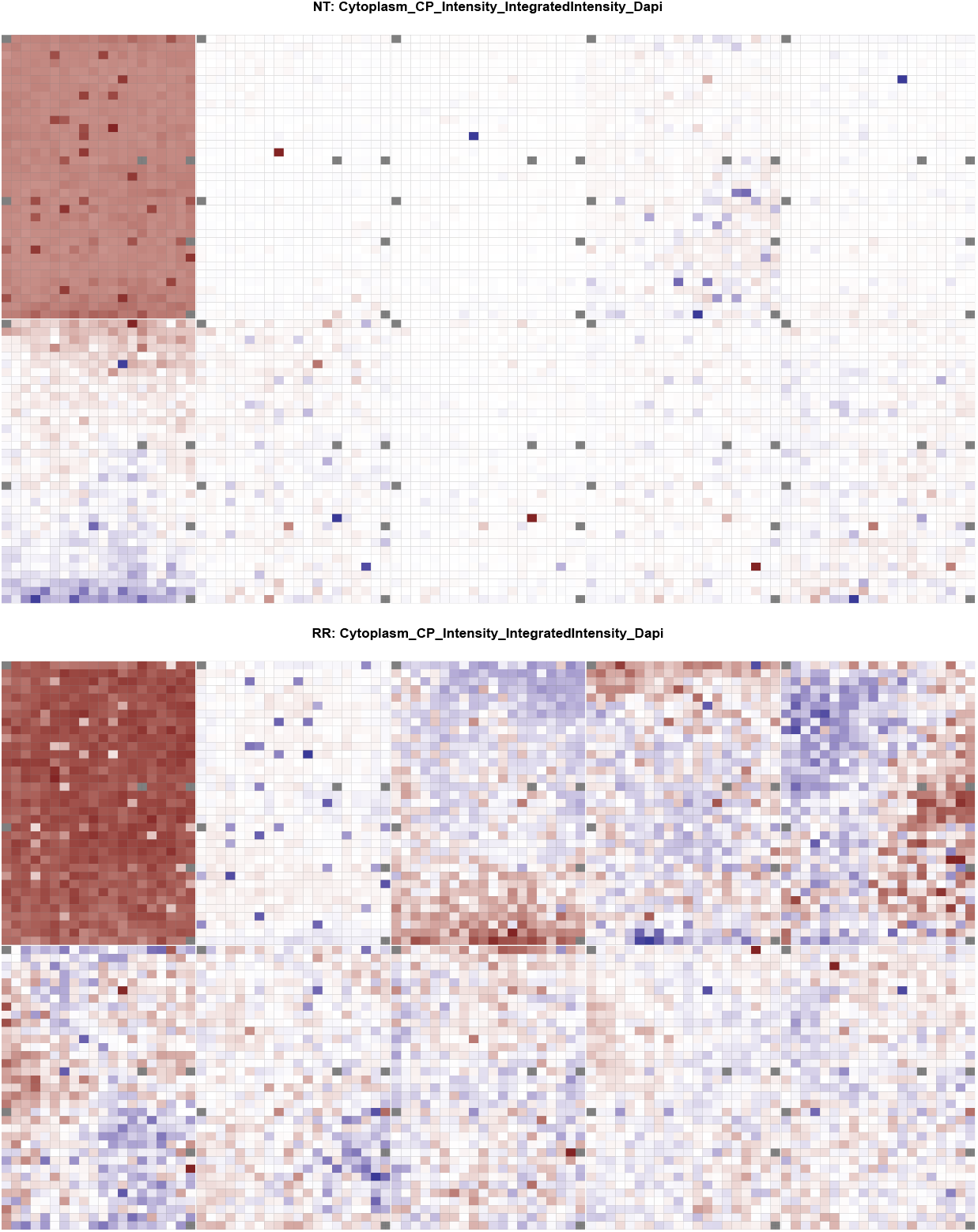
Heat map of elements of top ten right singular vectors for the total cytoplasmic DAPI intensity feature.

To see what biological effects can be found if we *a priori* remove the dominating ELN-NID effect, we re-analyze the MEMA data after removing these spots. Now we find an effect separating THBS from the other ECMps. This is particularly prominent in morphological features. As an example, in Figure 11 we plot a scatter plot of the top four RSVs for cell compactness. (RR) reveals a difference between THBS and the other ECMps. This effect shows up in many of the morphological features but not cell count. Thus this THBS effect does not appear to be of similar origin to the ELN-NID effect. Instead, it appears to be a biological effect on cell morphology.

**Fig. 11.**
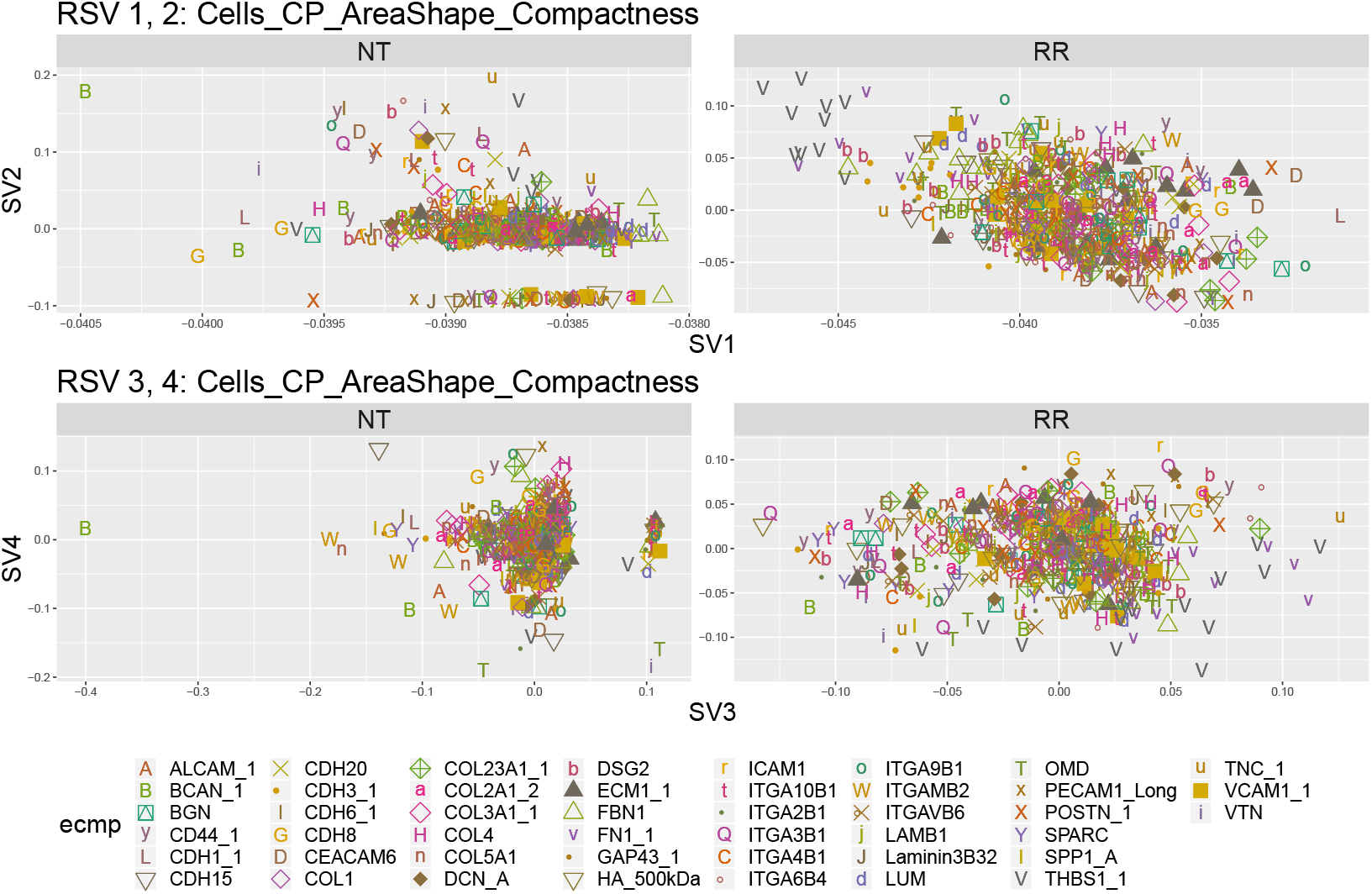
Scatter plot (after removing ELN and NID from analysis) of elements of top four right singular vectors against each other for the cell compactness feature. Shape and color indicate ECMp of the spot corresponding to the elements of the singular vector. The clusters seen in the NT panels are from missing spots on an outlier plate (see Supplementary Figure 31)

### 3.6. Data Integration for Discovering Within-Well Effects

In section 3.4 we saw that data integration helped make salient important between-well effects. In a similar fashion, the average right singular vectors (ASVs) help bring out within-well effects. In Figure 12 we plot the first two right ASVs against each other. Again, (RR) equitably integrates information from all features and helps highlight the NID/ELN effect.

**Fig. 12.**
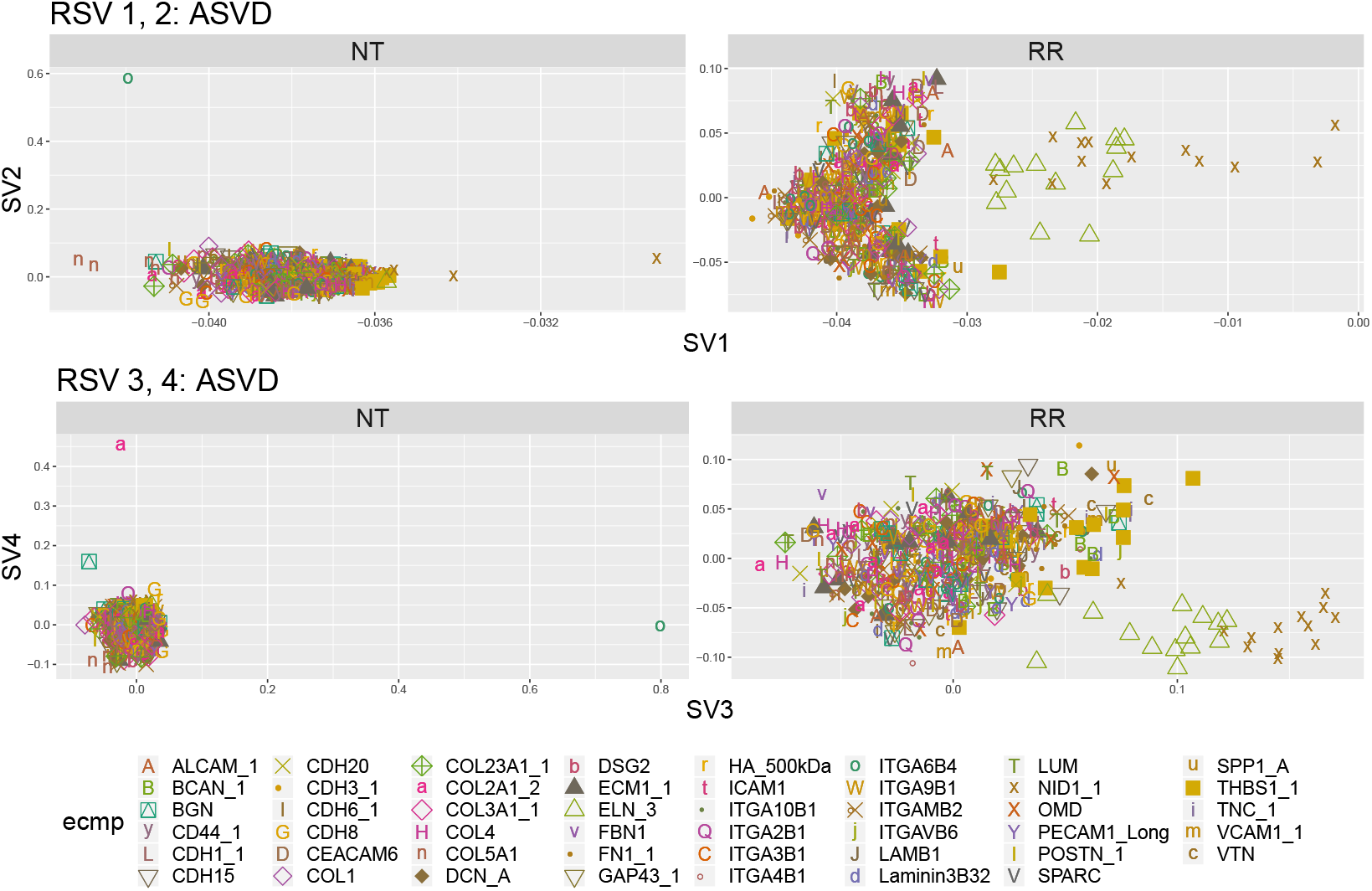
Scatter plot of elements of top four right ASVs calculated over 18 features measured on all MEMAs. Shape and color indicate ECMp of the spot corresponding to the elements of the singular vector.

Finally, we display heat-maps pseudo-images for the first ten right ASVs in Figure 13. The right ASVs for (NT) seem to be mostly picking up a couple of outliers. On the other hand, (RR) strongly picks up several interesting spatial effects.

**Fig. 13.**
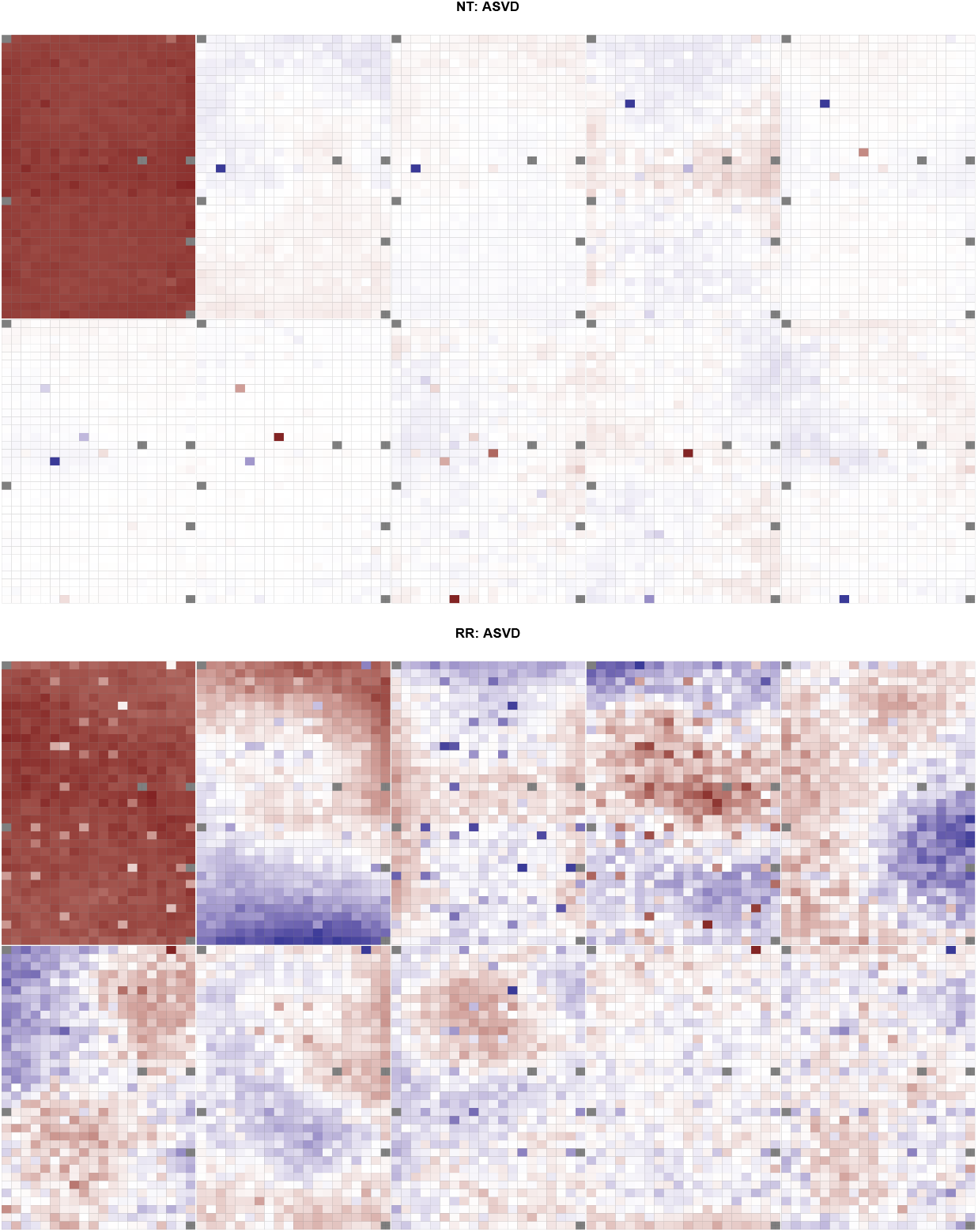
Heat-map of top ten right ASVs calculated over 18 features measured on all MEMAs.

## 4. Discussion

The microenvironment of cells is an important component of many cell and tissue level processes. Studying the cellular microenvironment not only furthers a fundamental understanding of these processes but also helps us understand the interaction of the microenvironment with disease-targeting therapies. In this paper we have explored the effects of several transformations as part of a pre-processing pipeline. The goal of these transformations is to emphasize important latent effects in the data and attenuate common and misleading aspects of MEMA data.

Un-transformed feature data is often encumbered by skewed measurement scales, outliers, or both. These aspects can hinder discovery of important biological effects and impair the identification of unwanted technical variation. To de-emphasize such misleading aspects of the data (O), (G), and their combination in (RR) were helpful. (O) removed outliers using a conservative threshold and (G) reduced skewness by Gaussianizing the data. Additionally, (RR) included a (Z) step that converted values to robust *z*-scores. (Z) and (RR) allowed features to be straight-forwardly integrated with a simple arithmetic average of Gram matrices.

We showed that a combination of a Gaussianizing transformation (G), *z*-score transformation (Z), and removal of outliers (O) can improve visualization and discovery of biological and technical latent effects in both features individually and when combining features together. Finally, as (RR) automatically chose transformations for each feature this allowed adaptive application of (RR) to a range of MEMA data containing different features.

This adaptive ability makes (RR) a promising candidate for pre-processing data generated by other image-based cell-profiling technologies. For example (RR) seems well suited for the analysis of data generated by Cyclic Immunofluorescence (Cy-cIF)(Lin *et al*., 2016). CycIF is a new technology that allows up to 30-channel immunofluorescent imaging. It does this through a series of imaging and washing steps, each step staining with up to 6 different stains. Thus, through image-analysis, CycIF has the potential to generate several hundred features. This is several times the number seen for MEMAs and further compounds the need for automatic feature transformation. Exploring the application of (RR) to other highly-multiplexed technologies like CycIf is a direction we hope to explore in future work.

## Supporting information

supplemental plots

## Notes

https://gjhunt.github.io/rr/

## References

Bhat, R. and Bissell, M. J. (2014). Of plasticity and specificity: dialectics of the microenvironment and macroenvironment and the organ phenotype. Wiley Interdisciplinary Reviews: Developmental Biology, 3(2), 147–163.

Bissell, M. J. and Labarge, M. A. (2005). Context, tissue plasticity, and cancer: are tumor stem cells also regulated by the microenvironment? Cancer cell, 7(1), 17–23.

Box, G. E. P. and Cox, D. R. (1964). An Analysis of Transformations. Journal of the Royal Statistical Society. Series B (Methodological), 26(2), 211–252.

Burger, J. A., Ghia, P., Rosenwald, A., and Caligaris-Cappio, F. (2009). The microenvironment in mature B-cell malignancies: a target for new treatment strategies. Blood, 114(16), 3367–75.

Caicedo, J. C., Cooper, S., Heigwer, F., Warchal, S., Qiu, P., Molnar, C., Vasilevich, A. S., Barry, J. D., Bansal, H. S., Kraus, O., Wawer, M., Paavolainen, L., Herrmann, M. D., Rohban, M., Hung, J., Hennig, H., Concannon, J., Smith, I., Clemons, P. A., Singh, S., Rees, P., Horvath, P., Linington, R. G., and Carpenter, A. E. (2017). Data-analysis strategies for image-based cell profiling. Nature Methods, 14(9), 849–863.

Gray, J., Heiser, L., and Korkola, J. (2014). Microenvironment Perturbagen (MEP) LINCS on Synapse.

Januschke, J. and Näthke, I. (2014). Stem cell decisions: A twist of fate or a niche market? Seminars in Cell & Developmental Biology, 34, 116–123.

Keenan, A. B., Jenkins, S. L., Jagodnik, K. M., Koplev, S., He, E., Torre, D., Wang, Z., Dohlman, A. B., Silverstein, M. C., Lachmann, A., Kuleshov, M. V., Ma’ayan, A., Stathias, V., Terryn, R., Cooper, D., Forlin, M., Koleti, A., Vidovic, D., Chung, C., Schürer, S. C., Vasiliauskas, J., Pilarczyk, M., Shamsaei, B., Fazel, M., Ren, Y., Niu, W., Clark, N. A., White, S., Mahi, N., Zhang, L., Kouril, M., Reichard, J. F., Sivaganesan, S., Medvedovic, M., Meller, J., Koch, R. J., Birtwistle, M. R., Iyengar, R., Sobie, E. A., Azeloglu, E. U., Kaye, J., Osterloh, J., Haston, K., Kalra, J., Finkbiener, S., Li, J., Milani, P., Adam, M., Escalante-Chong, R., Sachs, K., Lenail, A., Ramamoorthy, D., Fraenkel, E., Daigle, G., Hussain, U., Coye, A., Rothstein, J., Sareen, D., Ornelas, L., Banuelos, M., Mandefro, B., Ho, R., Svendsen, C. N., Lim, R. G., Stocksdale, J., Casale, M. S., Thompson, T. G., Wu, J., Thompson, L. M., Dardov, V., Venkatraman, V., Matlock, A., Van Eyk, J. E., Jaffe, J. D., Papanastasiou, M., Subramanian, A., Golub, T. R., Erickson, S. D., Fallahi-Sichani, M., Hafner, M., Gray, N. S., Lin, J. R., Mills, C. E., Muhlich, J. L., Niepel, M., Shamu, C. E., Williams, E. H., Wrobel, D., Sorger, P. K., Heiser, L. M., Gray, J. W., Korkola, J. E., Mills, G. B., LaBarge, M., Feiler, H. S., Dane, M. A., Bucher, E., Nederlof, M., Sudar, D., Gross, S., Kilburn, D. F., Smith, R., Devlin, K., Margolis, R., Derr, L., Lee, A., and Pillai, A. (2018). The Library of Integrated Network-Based Cellular Signatures NIH Program: System-Level Cataloging of Human Cells Response to Perturbations. Cell Systems, 6(1), 13–24.

LaBarge, M. (2013). Breaking the Canon: Indirect Regulation of Wnt Signaling in Mammary Stem Cells by MMP3. Cell Stem Cell, 13(3), 259–260.

LaBarge, M. A., Petersen, O. W., and Bissell, M. J. (2007). Of microenvironments and mammary stem cells. Stem cell reviews, 3(2), 137–46.

LaBarge, M. A., Nelson, C. M., Villadsen, R., Fridriksdottir, A., Ruth, J. R., Stampfer, M. R., Petersen, O. W., and Bissell, M. J. (2009). Human mammary progenitor cell fate decisions are products of interactions with combinatorial microenvironments. Integr. Biol., 1(1), 70–79.

Labarge, M. A., Parvin, B., and Lorens, J. B. (2014). Molecular deconstruction, detection, and computational prediction of microenvironment-modulated cellular responses to cancer therapeutics. Advanced drug delivery reviews, 69–70, 123-31.

Leek, J. T., Scharpf, R. B., Bravo, H. C., Simcha, D., Langmead, B., Johnson, W. E., Geman, D., Baggerly, K., and Irizarry, R. A. (2010). Tackling the widespread and critical impact of batch effects in high-throughput data. Nature Reviews Genetics, 11(10), 733–739.

Lin, C.-H., Lee, J. K., and LaBarge, M. A. (2012). Fabrication and Use of MicroEnvironment microArrays (MEArrays). Journal of Visualized Experiments, (68), 1–7.

Lin, C.-H., Jokela, T., Gray, J., and LaBarge, M. A. (2017). Combinatorial Microenvironments Impose a Continuum of Cellular Responses to a Single Pathway-Targeted Anti-cancer Compound. Cell Reports, 21(2), 533–545.

Lin, J.-R., Fallahi-Sichani, M., Chen, J.-Y., and Sorger, P. K. (2016). Cyclic Immunofluorescence (CycIF), A Highly Multiplexed Method for Single-cell Imaging. In Current Protocols in Chemical Biology, volume 8, pages 251–264. John Wiley & Sons, Inc., Hoboken, NJ, USA.

Maman, S. and Witz, I. P. (2018). A history of exploring cancer in context. Nature Reviews Cancer, 18(6), 359–376.

Pelissier, F. A., Garbe, J. C., Ananthanarayanan, B., Miyano, M., Lin, C. H., Jokela, T., Kumar, S., Stampfer, M. R., Lorens, J. B., and LaBarge, M. A. (2014). Age-Related Dysfunction in Mechanotransduction Impairs Differentiation of Human Mammary Epithelial Progenitors. Cell Reports, 7(6), 1926–1939.

R Core Team (2018). R: A Language and Environment for Statistical Computing. R Foundation for Statistical Computing, Vienna, Austria.

Smith, R., Devlin, K., Kilburn, D., Gross, S., Sudar, D., Bucher, E., Nederlof, M., Dane, M., Gray, J. W., Heiser, L., and Korkola, J. E. (2019). Using Microarrays to Interrogate Microenvironmental Impact on Cellular Phenotypes in Cancer. Journal of visualized experiments: JoVE.

Teti, A. (1992). Regulation of Cellular Functions by Extracellular Matrix. Technical report.

Watson, S. S., Dane, M., Chin, K., Jonas, O., Gray, J. W., and Korkola, J. E. (2018). Microenvironment-Mediated Mechanisms of Resistance to HER2 Inhibitors Differ between HER2+ Breast Cancer Subtypes. Cell Systems, 6, 329–342.e6.

