## supplemental plots for "Transformation and Integration of Microenvironment Microarray Data Improves Discovery of Latent Effects"

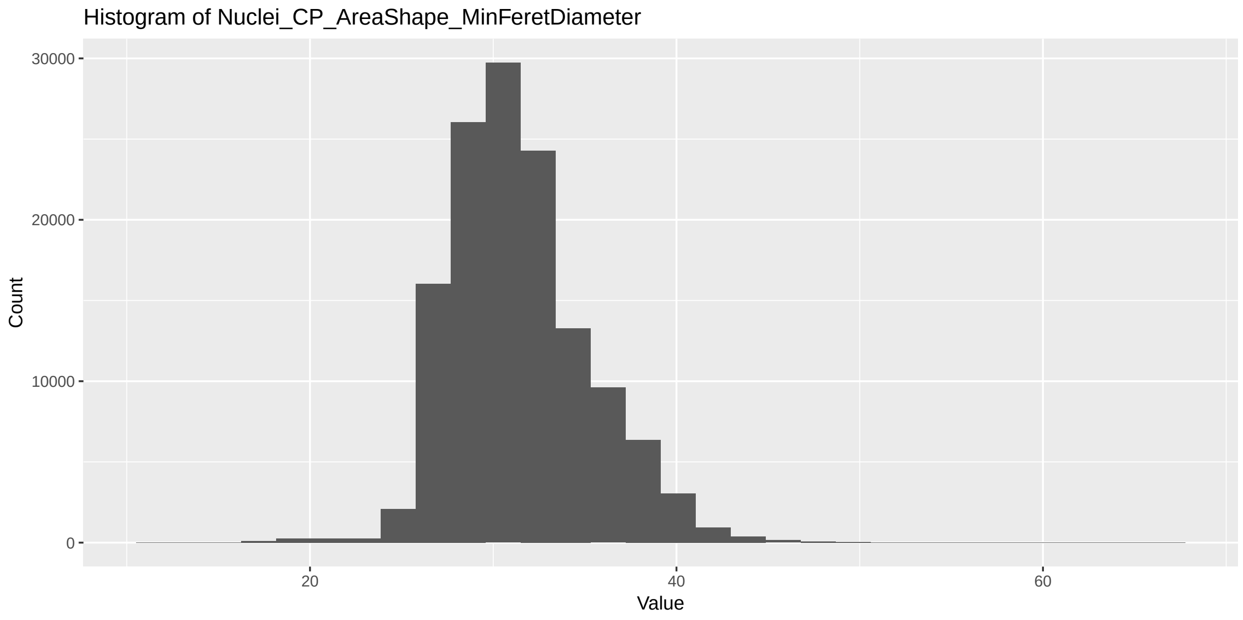


Figure 1: Histogram of nuclei orientation across all wells and spots.

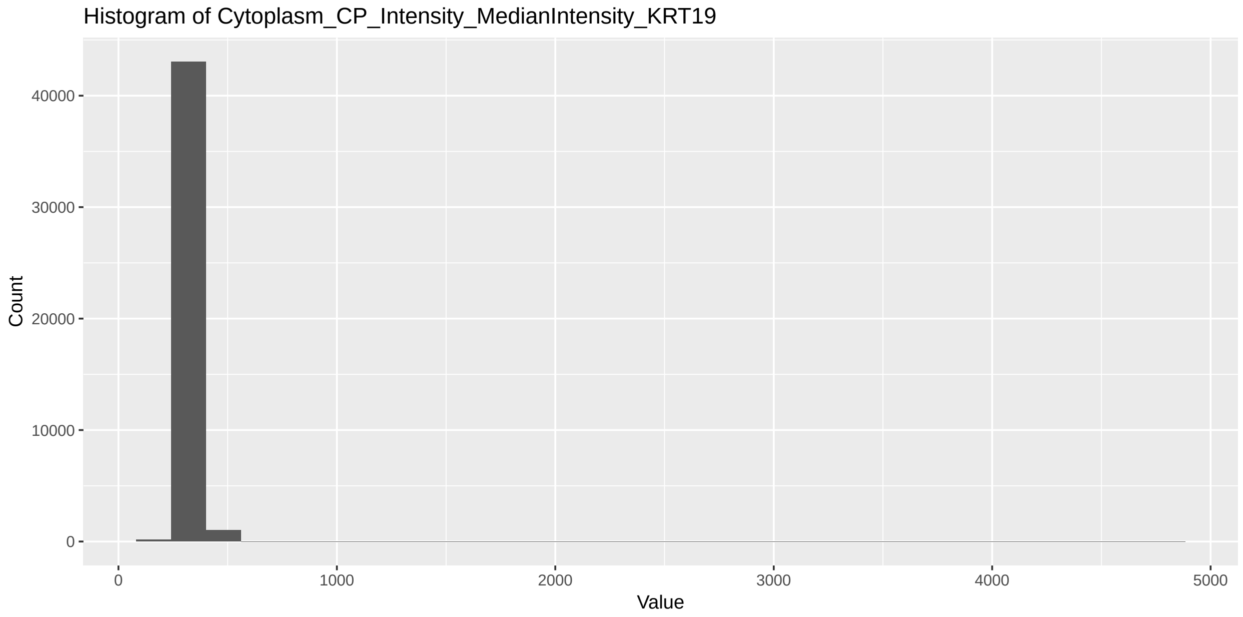


Figure 2: Histogram of total nuclei DAPI intensity across all wells and spots.

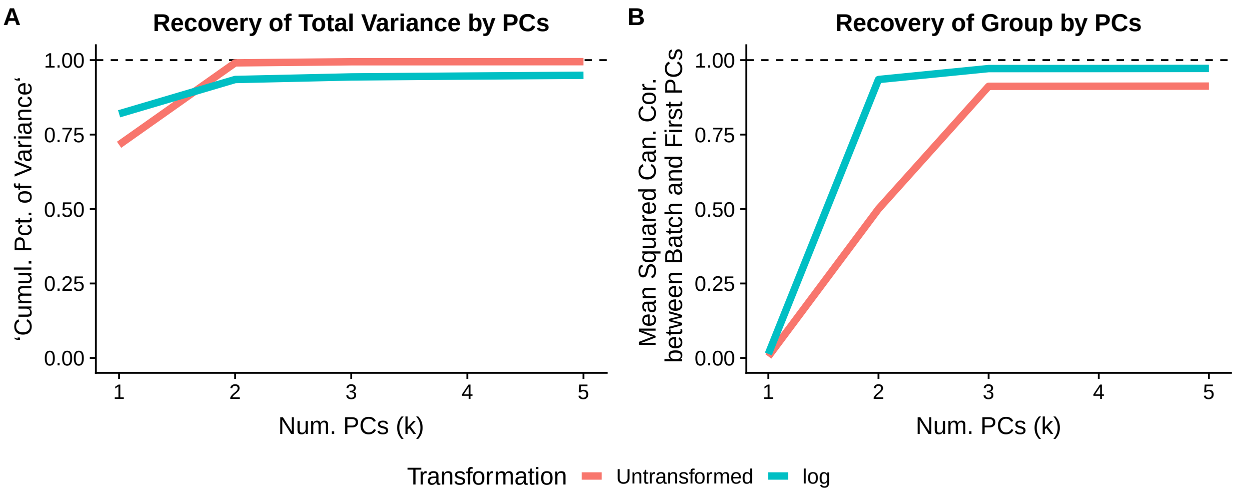


Figure 3: (A) The percentage of cumulative variance captured by first $k$ principal components for both un-transformed data and log-transformed data. (B) The mean squared canonical correlations between the grouping factor and the first $k$ principal components.

For clarity, we reproduce the explanation of (B) from main text.

We assess the transformations by measuring the percentage of the batch (group difference) captured by the first $k$ singular vectors of the transformed feature matrix. Let $U = [u_1, \ldots, u_N] \in \mathbb{R}^{M \times N}$ be the (complete) left singular vectors of a feature matrix and $B \in \mathbb{R}^{M \times D}$ be the batch indicator matrix so that $B_{ij} = 1$ if well $i$ is in batch $j$ for $j = 1, \ldots, D$. Here we have $D = 2$ for the two groups. For $k = 1, \ldots, N$ and $t = 1 \ldots \min(k, D)$ define $C_k^{(t)}$ to be the $t^{th}$ canonical correlation between the first $k$ left singular vectors $U_k = [u_1, \ldots, u_k]$ and the batch $B$. Then let

$$C_k^2 = \frac{1}{D} \sum_{t=1}^{\min(k,D)} \left( C_k^{(t)} \right)^2$$

to be the average of these squared canonical correlations. We can interpret $C_k^2$ as the percentage of the batch $B$ that is captured by these first $k$ singular vectors.

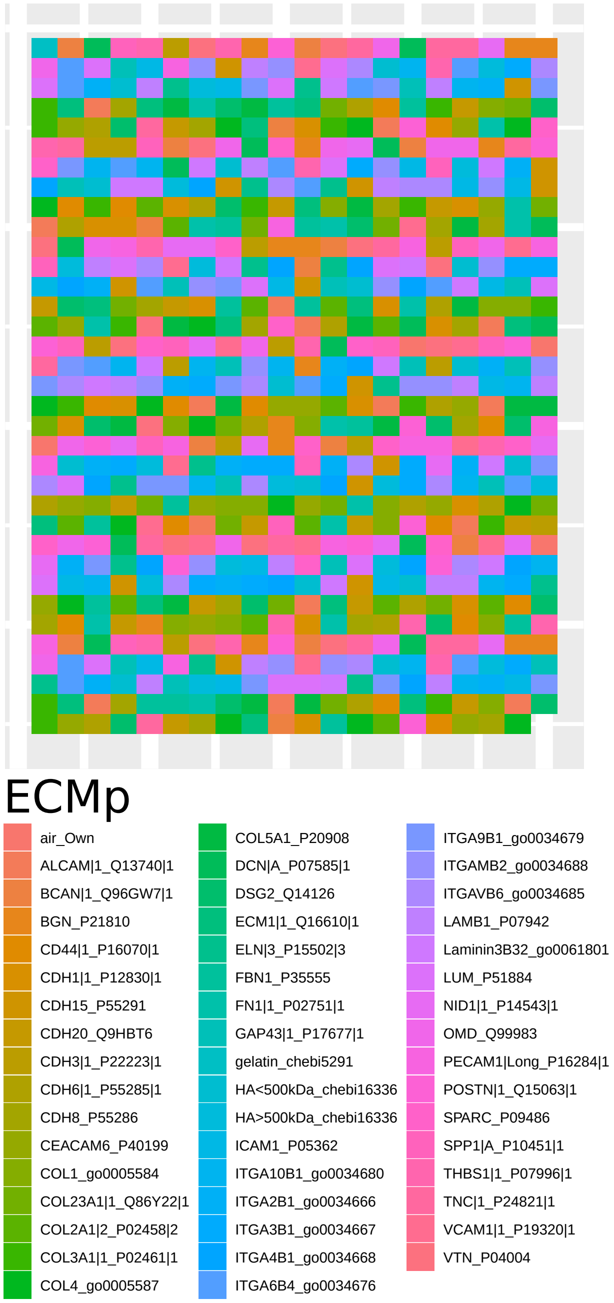


Figure 4: MEMA layout design of ECMps for each well. ECMp layout is identical across all wells.

| Feature |
|---|
| Cells_CP_AreaShape_Area |
| Cells_CP_AreaShape_Compactness |
| Cells_CP_AreaShape_Eccentricity |
| Cells_CP_AreaShape_Extent |
| Cells_CP_AreaShape_FormFactor |
| Cells_CP_AreaShape_MajorAxisLength |
| Cells_CP_AreaShape_MaxFeretDiameter |
| Cells_CP_AreaShape_MaximumRadius |
| Cells_CP_AreaShape_MeanRadius |
| Cells_CP_AreaShape_MedianRadius |
| Cells_CP_AreaShape_MinFeretDiameter |
| Cells_CP_AreaShape_MinorAxisLength |
| Cells_CP_AreaShape_Perimeter |
| Cells_CP_AreaShape_Solidity |
| Cytoplasm_CP_AreaShape_Area |
| Cytoplasm_CP_AreaShape_Compactness |
| Cytoplasm_CP_AreaShape_Eccentricity |
| Cytoplasm_CP_AreaShape_Extent |
| Cytoplasm_CP_AreaShape_FormFactor |
| Cytoplasm_CP_AreaShape_MajorAxisLength |
| Cytoplasm_CP_AreaShape_MaxFeretDiameter |
| Cytoplasm_CP_AreaShape_MaximumRadius |
| Cytoplasm_CP_AreaShape_MeanRadius |
| Cytoplasm_CP_AreaShape_MedianRadius |
| Cytoplasm_CP_AreaShape_MinFeretDiameter |
| Cytoplasm_CP_AreaShape_MinorAxisLength |
| Cytoplasm_CP_AreaShape_Perimeter |
| Cytoplasm_CP_AreaShape_Solidity |
| Nuclei_CP_AreaShape_Area |
| Nuclei_CP_AreaShape_Compactness |
| Nuclei_CP_AreaShape_Eccentricity |
| Nuclei_CP_AreaShape_Extent |
| Nuclei_CP_AreaShape_FormFactor |
| Nuclei_CP_AreaShape_MajorAxisLength |
| Nuclei_CP_AreaShape_MaxFeretDiameter |
| Nuclei_CP_AreaShape_MaximumRadius |
| Nuclei_CP_AreaShape_MeanRadius |
| Nuclei_CP_AreaShape_MedianRadius |
| Nuclei_CP_AreaShape_MinFeretDiameter |
| Nuclei_CP_AreaShape_MinorAxisLength |
| Nuclei_CP_AreaShape_Orientation |
| Nuclei_CP_AreaShape_Perimeter |
| Nuclei_CP_AreaShape_Solidity |
| Cells_CP_Intensity_IntegratedIntensity_CellMask |
| Cells_CP_Intensity_IntegratedIntensity_KRT19 |
| Cells_CP_Intensity_IntegratedIntensity_KRT5 |
| Cells_CP_Intensity_MedianIntensity_CellMask |
| Cells_CP_Intensity_MedianIntensity_KRT19 |
| Cells_CP_Intensity_MedianIntensity_KRT5 |
| Cytoplasm_CP_Intensity_IntegratedIntensity_CellMask |
| Cytoplasm_CP_Intensity_IntegratedIntensity_Dapi |
| Cytoplasm_CP_Intensity_IntegratedIntensity_KRT19 |
| Cytoplasm_CP_Intensity_IntegratedIntensity_KRT5 |
| Cytoplasm_CP_Intensity_MedianIntensity_CellMask |
| Cytoplasm_CP_Intensity_MedianIntensity_Dapi |
| Cytoplasm_CP_Intensity_MedianIntensity_KRT19 |
| Cytoplasm_CP_Intensity_MedianIntensity_KRT5 |
| Nuclei_CP_Intensity_IntegratedIntensity_Dapi |
| Nuclei_CP_Intensity_IntegratedIntensity_KRT19 |
| Nuclei_CP_Intensity_IntegratedIntensity_KRT5 |
| Nuclei_CP_Intensity_MedianIntensity_Dapi |
| Nuclei_CP_Intensity_MedianIntensity_KRT19 |
| Nuclei_CP_Intensity_MedianIntensity_KRT5 |
| Cytoplasm_PA_Intensity_LineageRatio |
| Spot_PA_SpotCellCount |
| Cells_CP_Intensity_IntegratedIntensity_Actin |
| Cells_CP_Intensity_IntegratedIntensity_MitoTracker |
| Cells_CP_Intensity_MedianIntensity_Actin |
| Cells_CP_Intensity_MedianIntensity_MitoTracker |
| Cytoplasm_CP_Intensity_IntegratedIntensity_Actin |
| Cytoplasm_CP_Intensity_IntegratedIntensity_MitoTracker |
| Cytoplasm_CP_Intensity_MedianIntensity_Actin |
| Cytoplasm_CP_Intensity_MedianIntensity_MitoTracker |
| Nuclei_CP_Texture_AngularSecondMoment_Fibrillarin_3_0 |
| Nuclei_CP_Texture_AngularSecondMoment_Fibrillarin_3_90 |
| Nuclei_CP_Texture_Contrast_Fibrillarin_3_0 |
| Nuclei_CP_Texture_Contrast_Fibrillarin_3_90 |
| Nuclei_CP_Texture_Correlation_Fibrillarin_3_0 |
| Nuclei_CP_Texture_Correlation_Fibrillarin_3_90 |
| Nuclei_CP_Texture_DifferenceEntropy_Fibrillarin_3_0 |
| Nuclei_CP_Texture_DifferenceEntropy_Fibrillarin_3_90 |
| Nuclei_CP_Texture_DifferenceVariance_Fibrillarin_3_0 |
| Nuclei_CP_Texture_DifferenceVariance_Fibrillarin_3_90 |
| Nuclei_CP_Texture_Entropy_Fibrillarin_3_0 |
| Nuclei_CP_Texture_Entropy_Fibrillarin_3_90 |
| Nuclei_CP_Texture_InfoMeas1_Fibrillarin_3_0 |
| Nuclei_CP_Texture_InfoMeas1_Fibrillarin_3_90 |
| Nuclei_CP_Texture_InfoMeas2_Fibrillarin_3_0 |
| Nuclei_CP_Texture_InfoMeas2_Fibrillarin_3_90 |
| Nuclei_CP_Texture_InverseDifferenceMoment_Fibrillarin_3_0 |
| Nuclei_CP_Texture_InverseDifferenceMoment_Fibrillarin_3_90 |
| Nuclei_CP_Texture_SumAverage_Fibrillarin_3_0 |
| Nuclei_CP_Texture_SumAverage_Fibrillarin_3_90 |
| Nuclei_CP_Texture_SumEntropy_Fibrillarin_3_0 |
| Nuclei_CP_Texture_SumEntropy_Fibrillarin_3_90 |
| Nuclei_CP_Texture_SumVariance_Fibrillarin_3_0 |
| Nuclei_CP_Texture_SumVariance_Fibrillarin_3_90 |
| Nuclei_CP_Texture_Variance_Fibrillarin_3_0 |
| Nuclei_CP_Texture_Variance_Fibrillarin_3_90 |
| Nuclei_CP_Intensity_IntegratedIntensity_EdU |
| Nuclei_CP_Intensity_IntegratedIntensity_Fibrillarin |
| Nuclei_CP_Intensity_MedianIntensity_EdU |
| Nuclei_CP_Intensity_MedianIntensity_Fibrillarin |

Table 1: List of all features extracted from at least one MEMA plate.

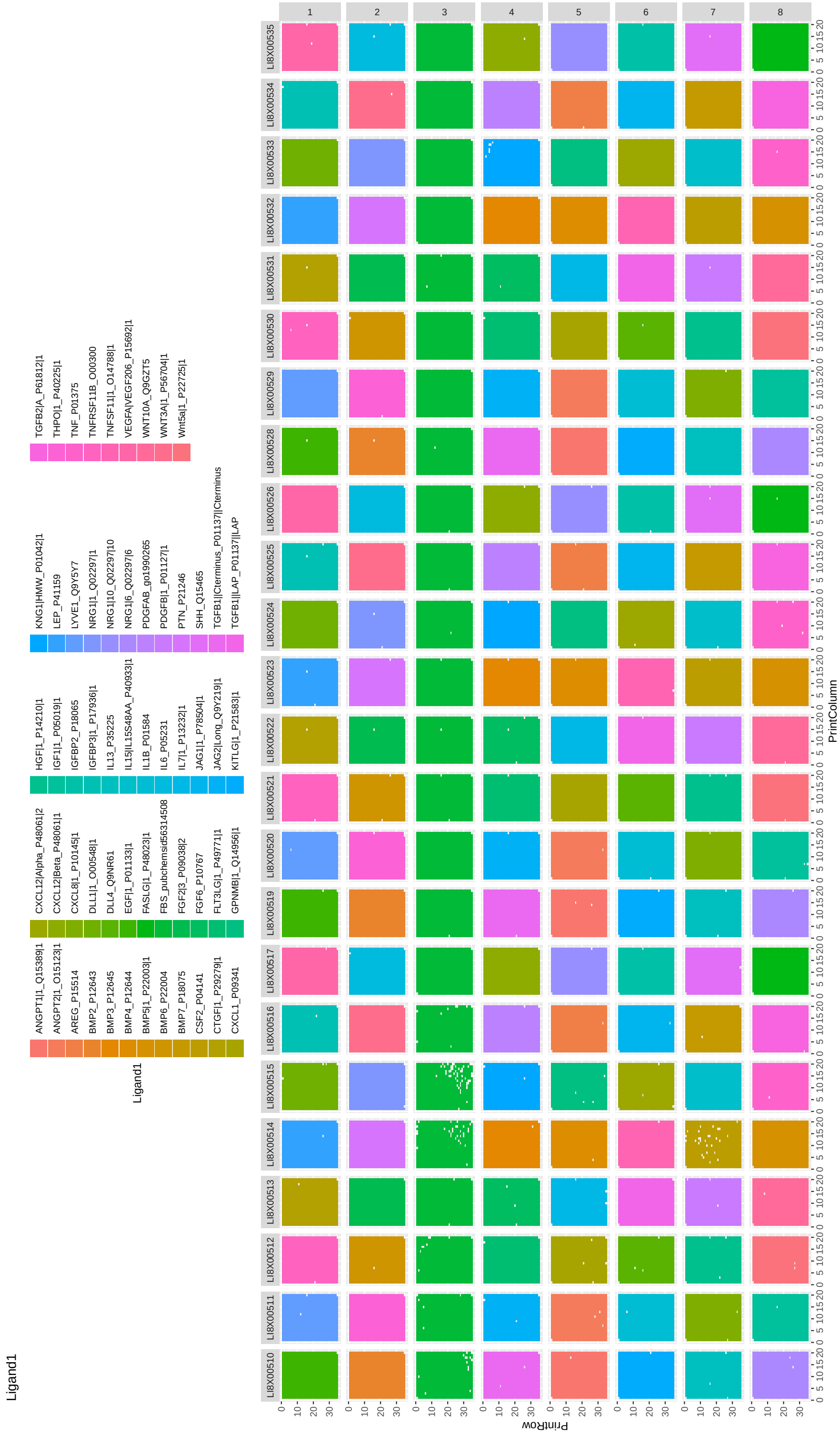


Figure 5: MEMA layout of ligands across plates and wells. Each row is a plate consisting of eight wells (columns). Color indicates ligand added to buffer solution of the well.

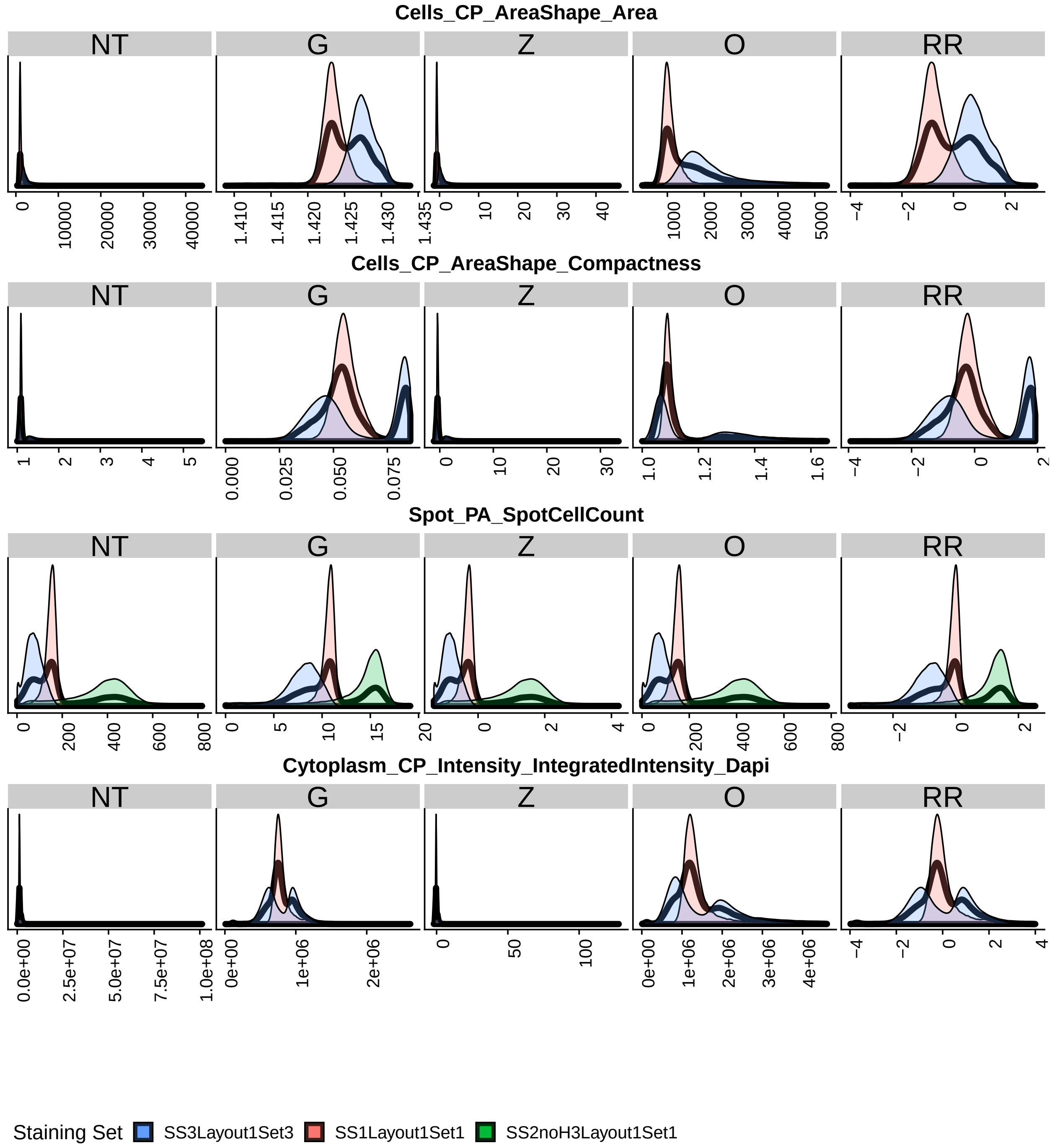


Figure 6: Density of elements of feature matrices. Black density is all elements combined. Colored densities are the densities denote staining batch. Subplots are for five processing transformations of this matrix: (NT) no transformation, (G) Gaussianization, (Z) $z$-score, (O) outlier removal, (RR) the three-step (G), (Z), and (O), robust re-scaling.

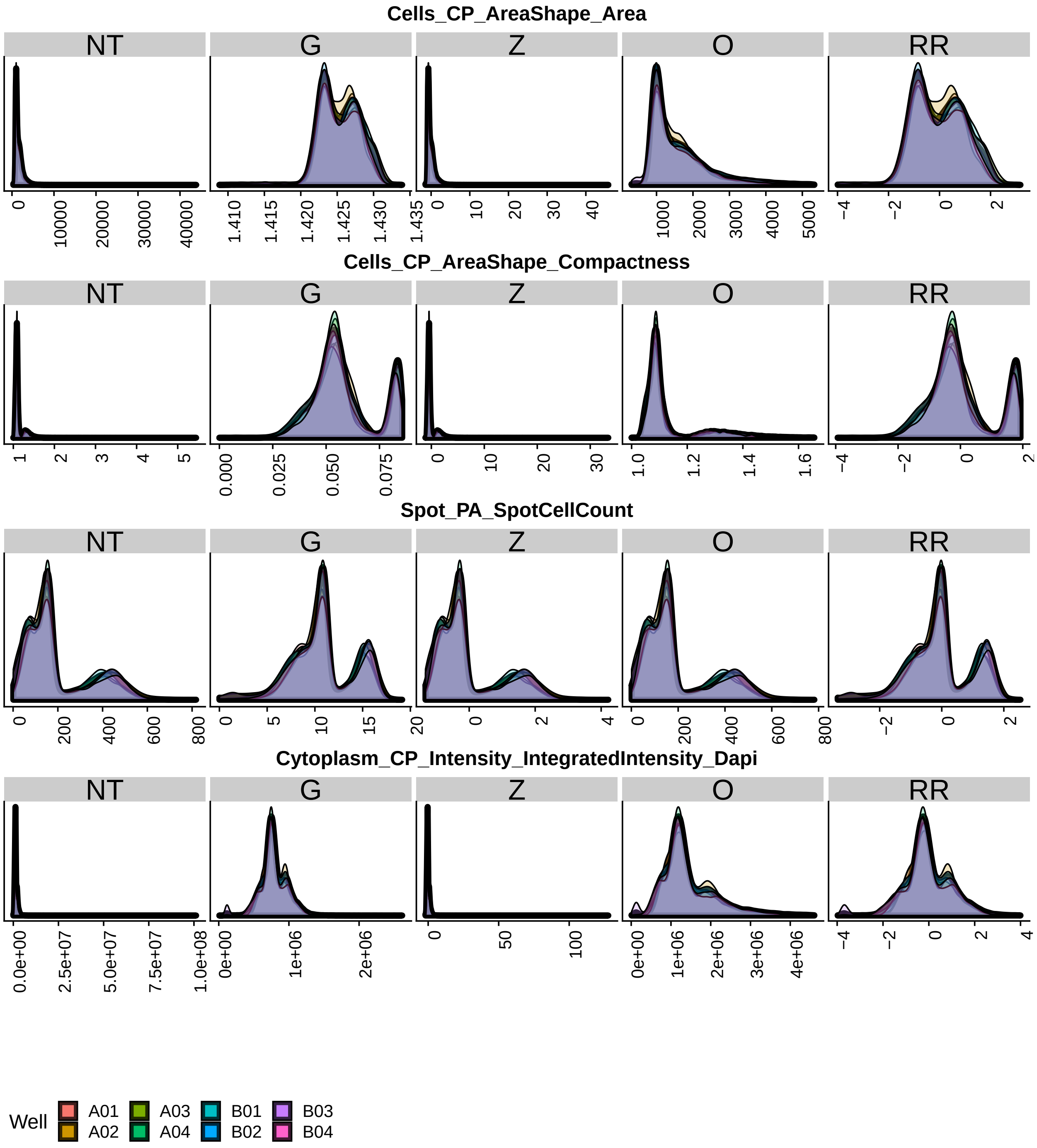


Figure 7: Similar to Figure 6 except colors indicate well.

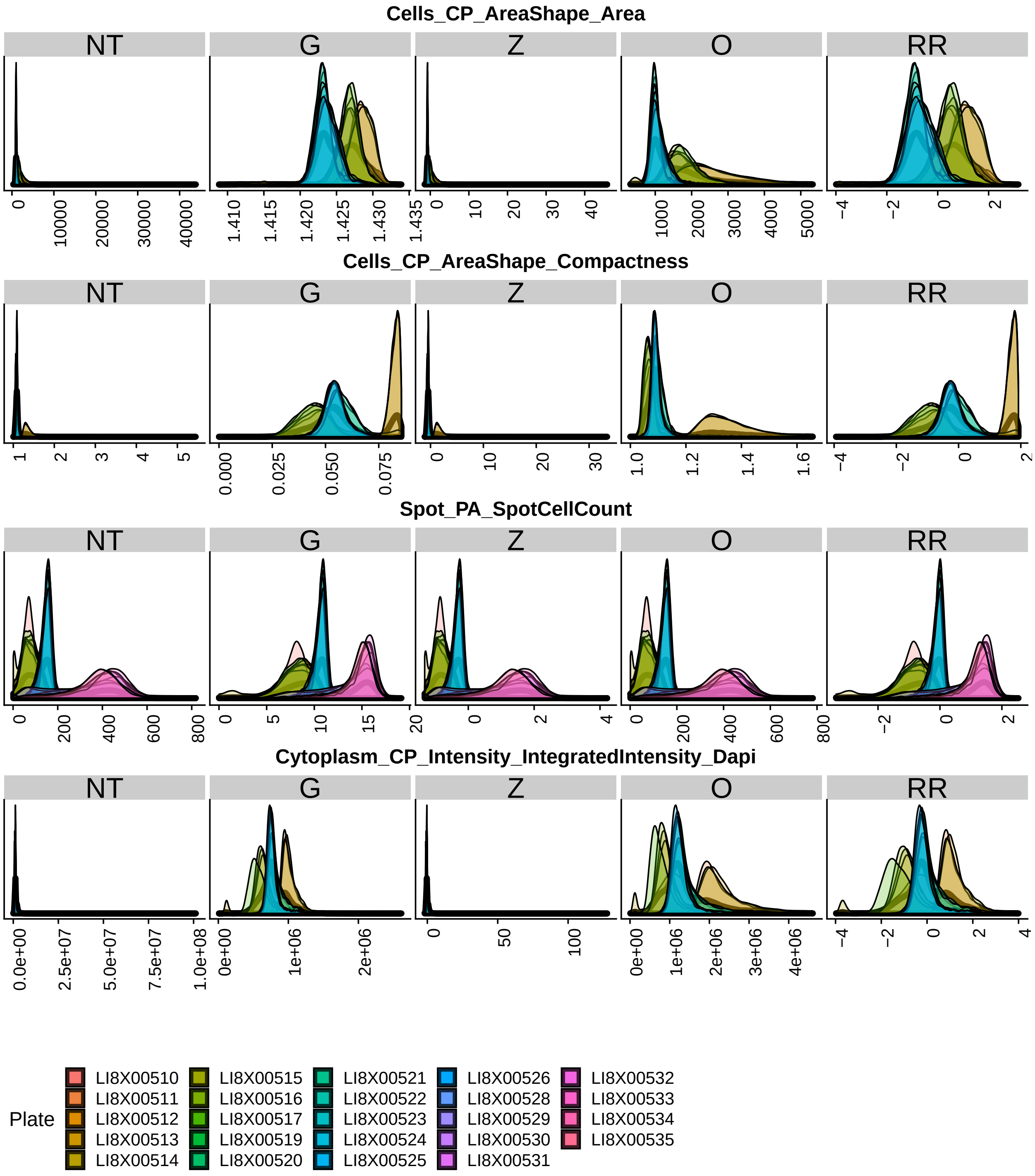


Figure 8: Similar to Figure 6 except colors indicate plate.

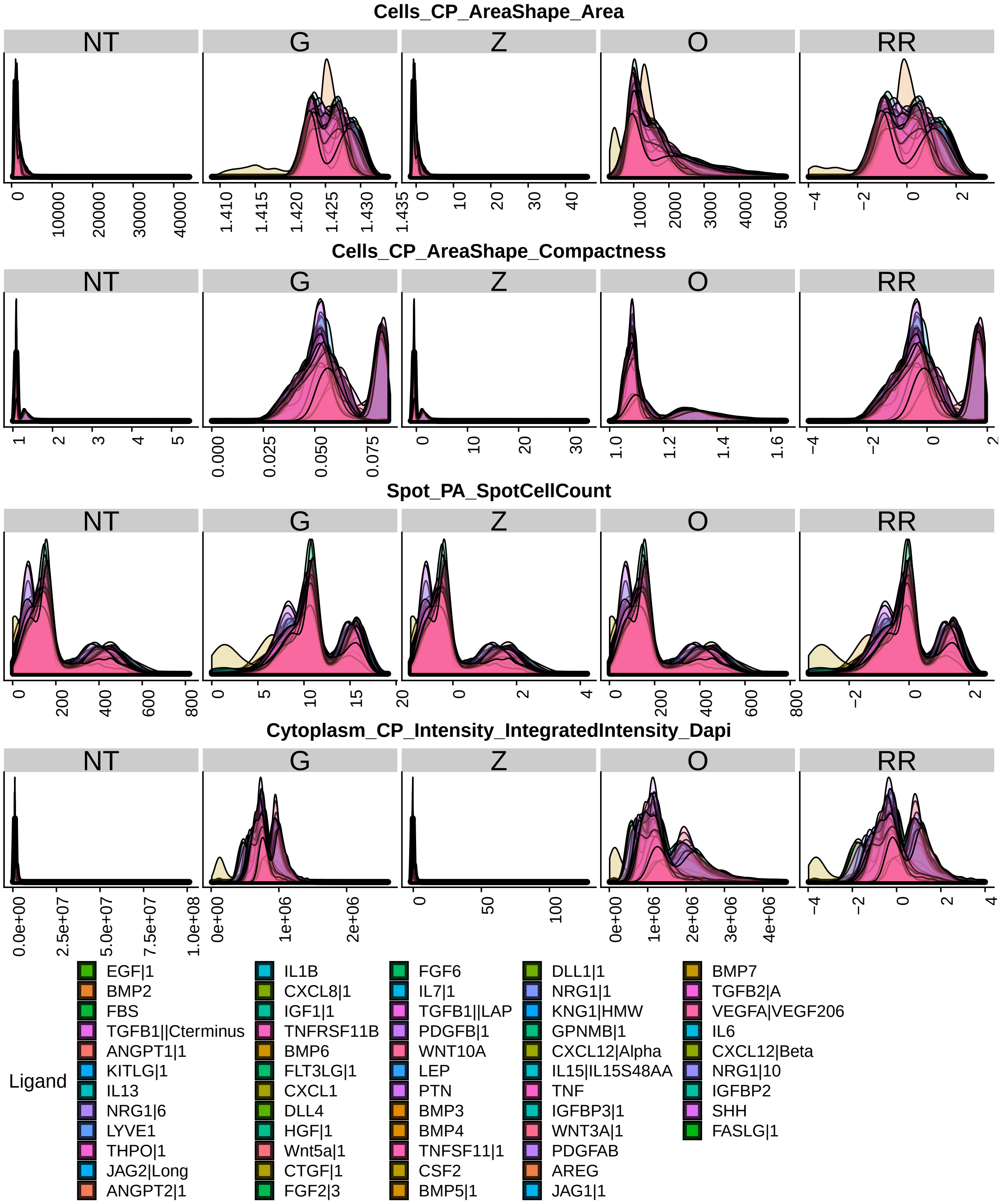


Figure 9: Similar to Figure 6 except colors indicate ligand.

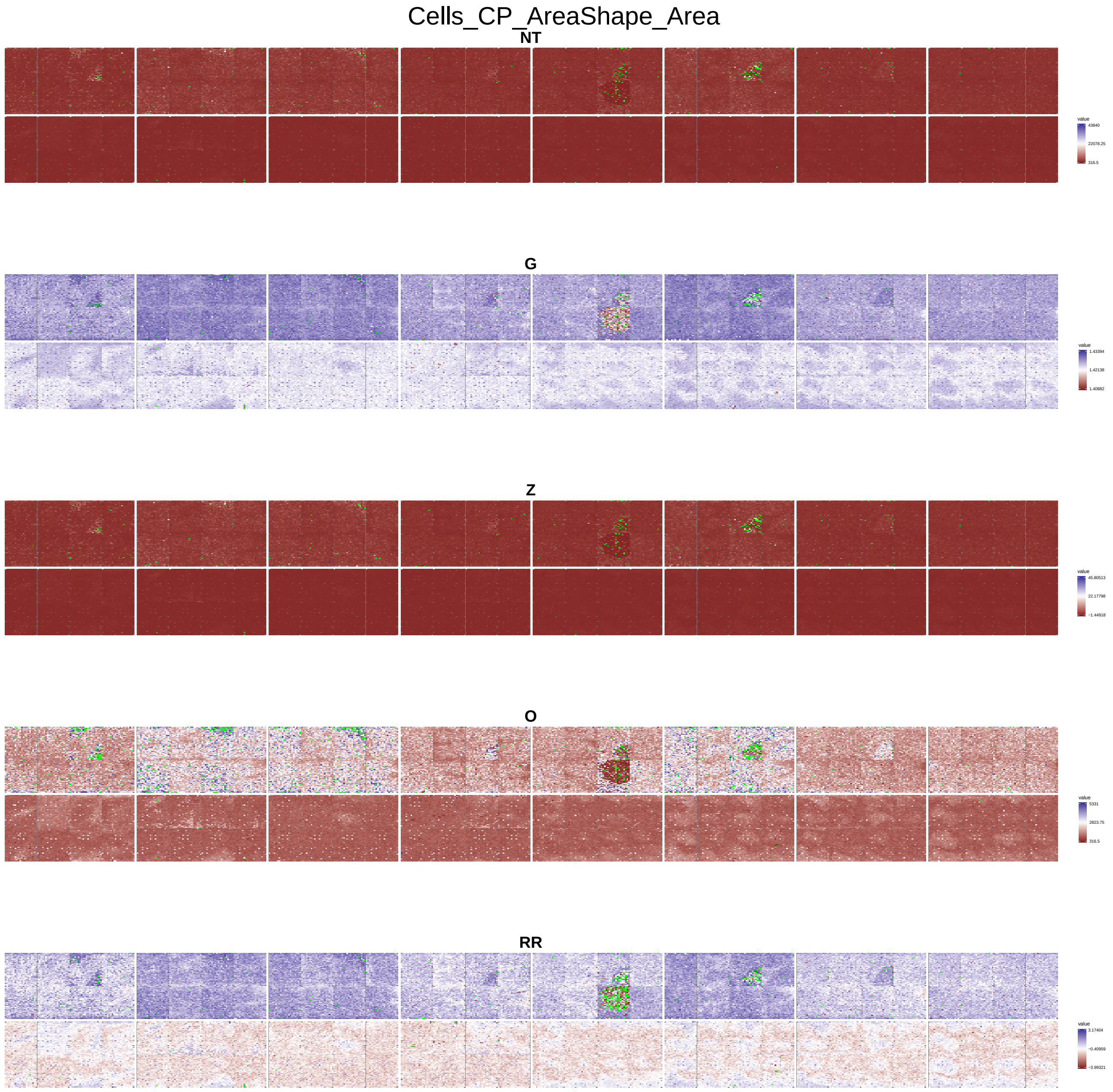


Figure 10: The next series of plots are heat-maps of MEMA plates across the five transformations (NT), (G), (Z), (O), (RR). Rows of each plot are the staining three batches. Colors are more blue if they are close to the minimum, red if they are close to the maximum, and white if they are close to half-way between. Green spots are missing. Dark grey spots are omitted according to the MEMA design.

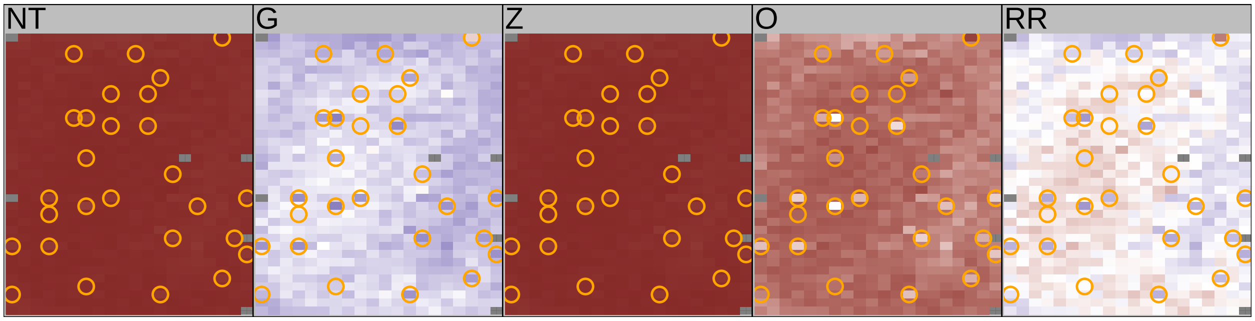


Figure 11: Heat map of a single well across the five transformations (NT), (G), (Z), (O), (RR). This is a sub-plot of Supplementary Figure 10. Color scaled is determined globally over all spots, wells, and plates in the dataset to reflect the fact that the transformation is similarly calculated over this data. Thus we see no blue in this (NT) sub-plot as we see almost no blue in Supplementary Figure 10. This plot is a representative microcosm of the larger plot. Orange circles highlight the ELN and NID1 spots.

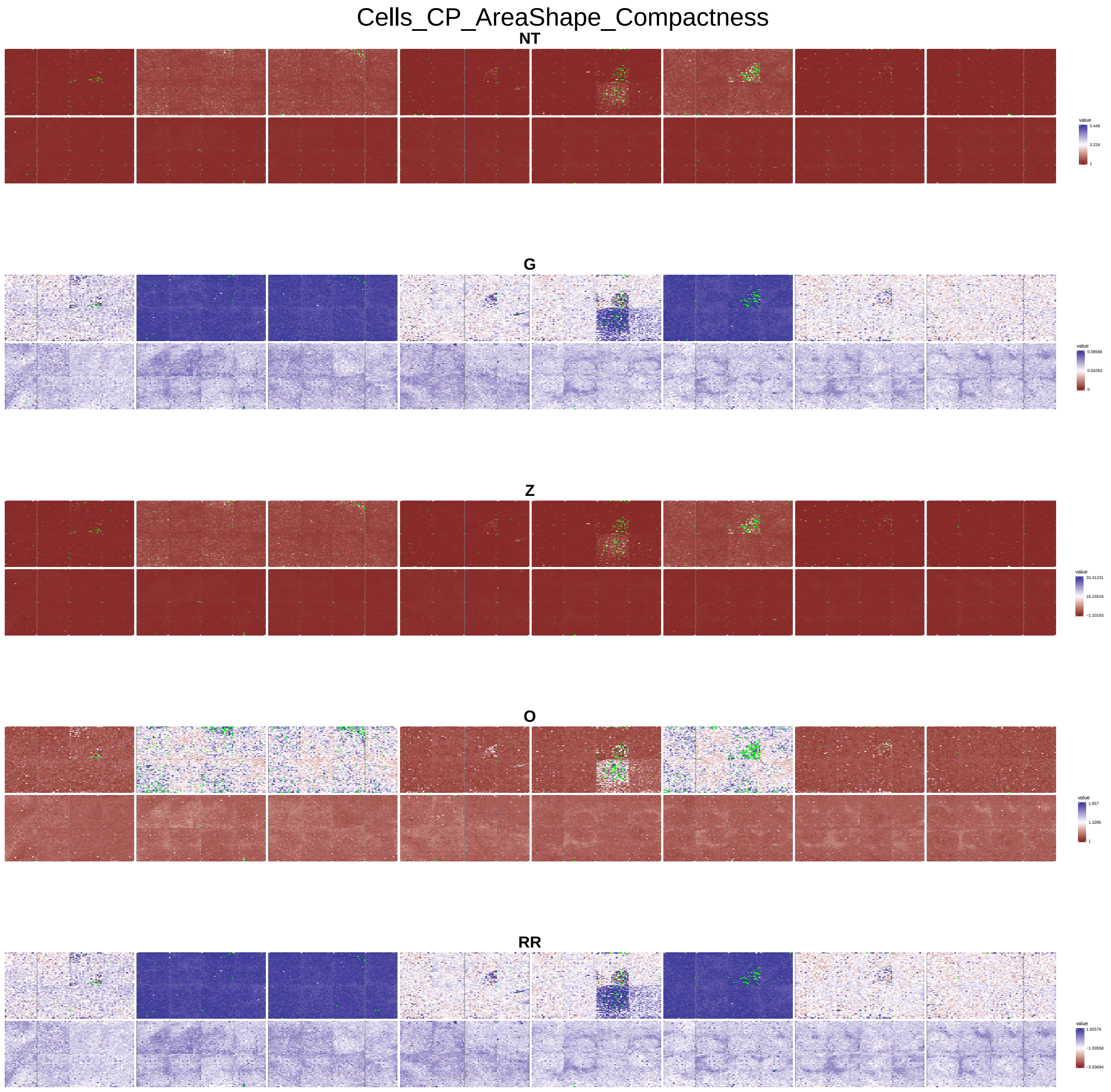


Figure 12: Similar to Figure 10 but for compactness.

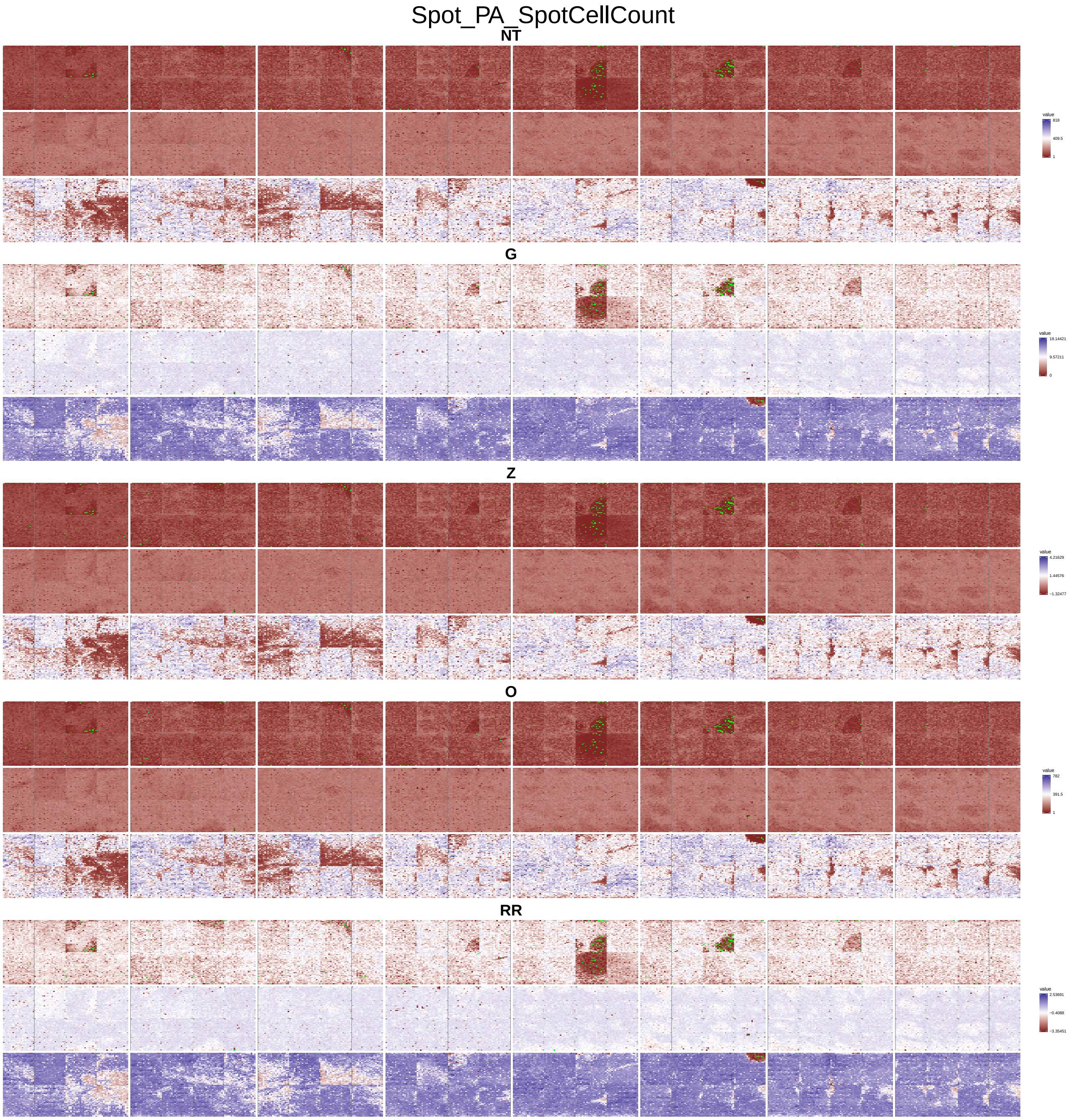


Figure 13: Similar to Figure 10 but for cell count.

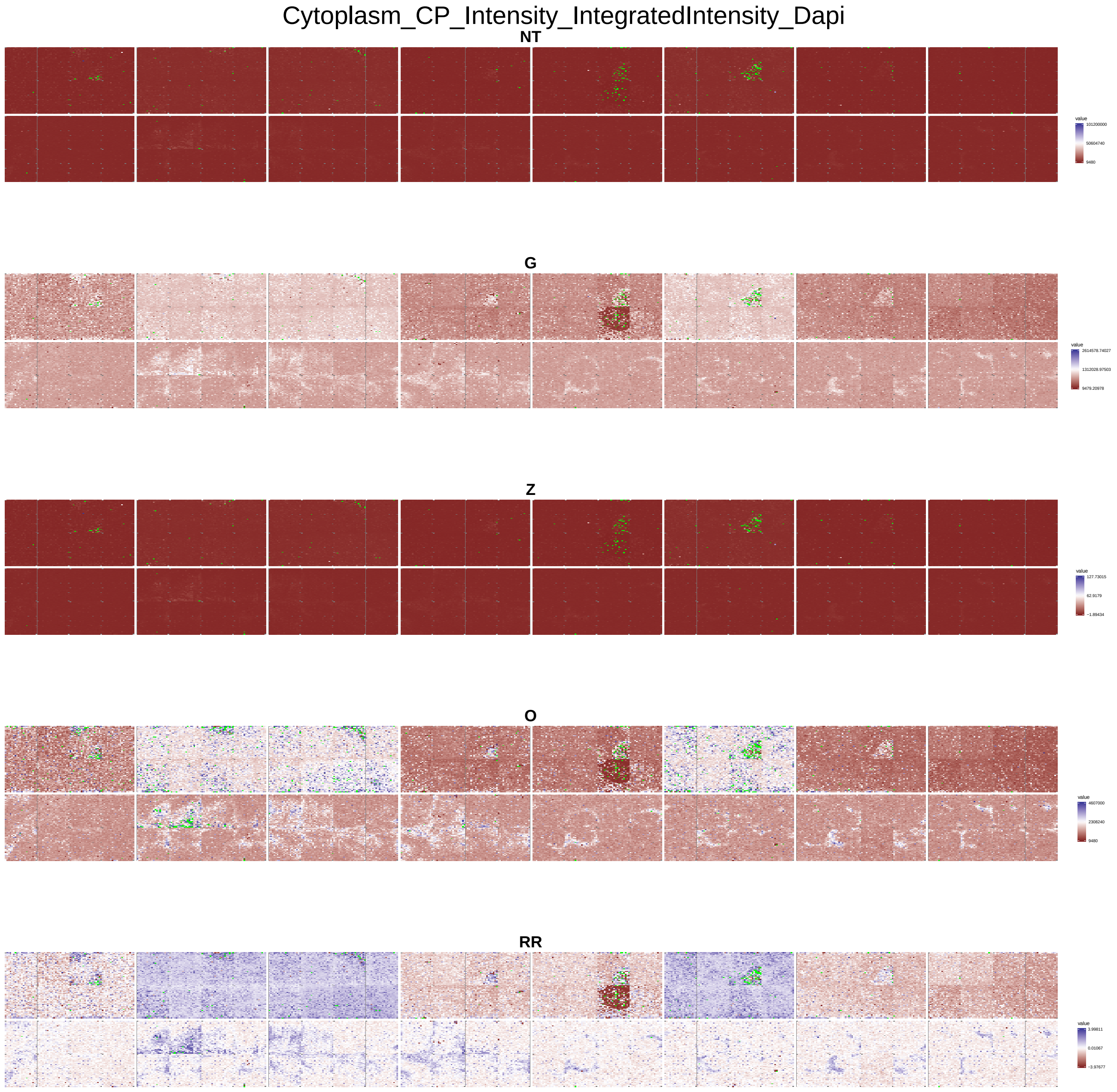


Figure 14: Similar to Figure 10 for for DAPI intensity.

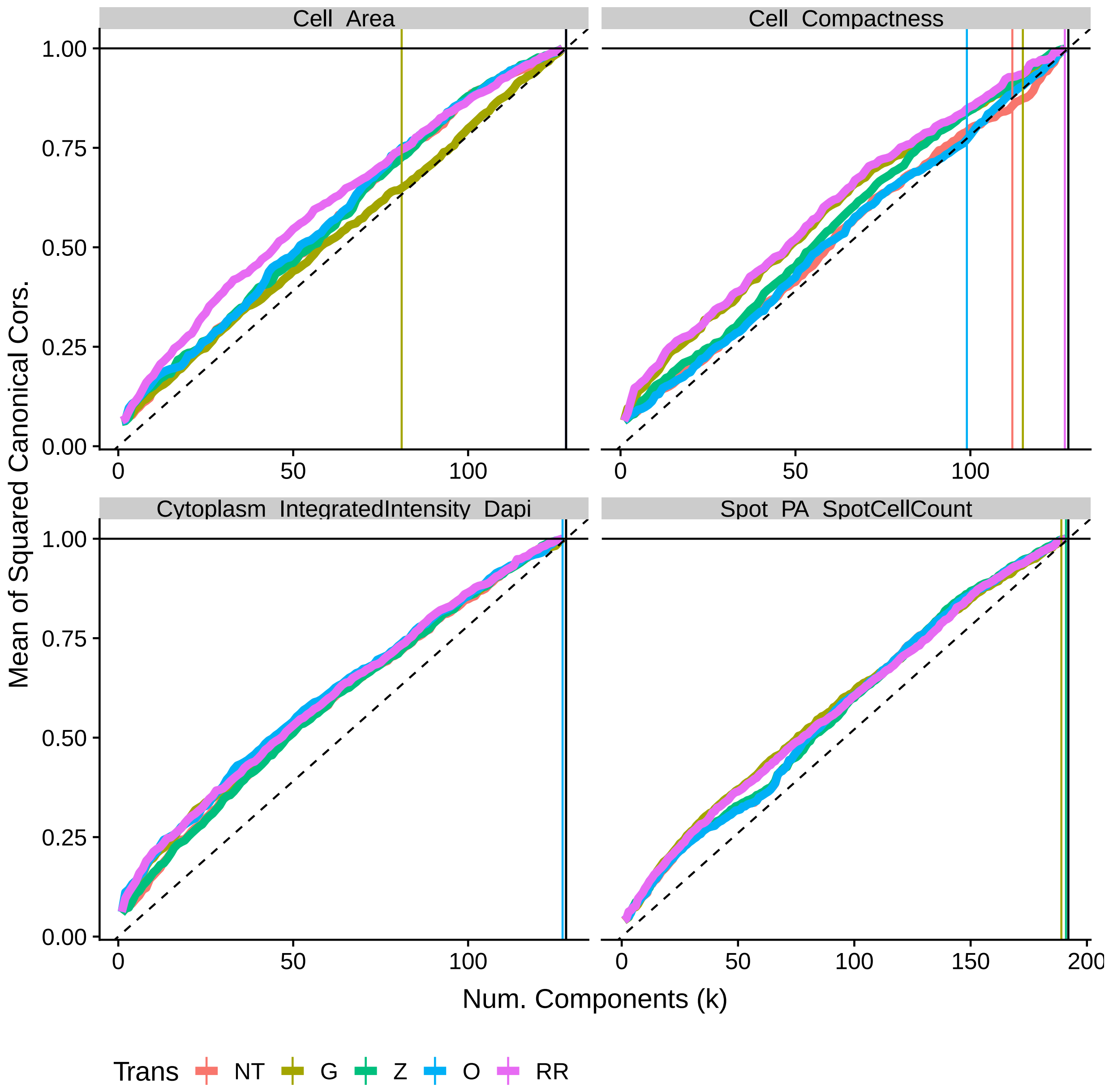


Figure 15: Mean of the squared canonical correlations between the first $k$ principal components and the plate batch indicator variables.

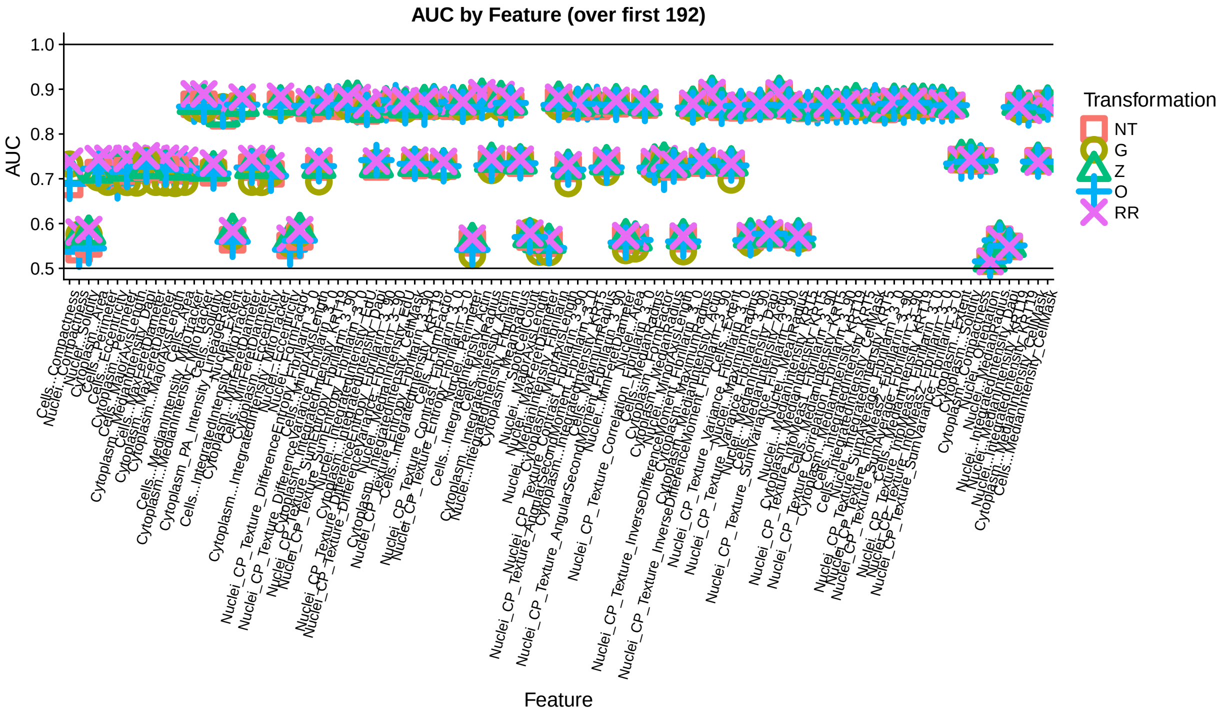


Figure 16: Grand mean of the squared canonical correlations across number of components ($k$). Canonical correlation is calculated between the first $k$ principal components and the plate indicator variables.

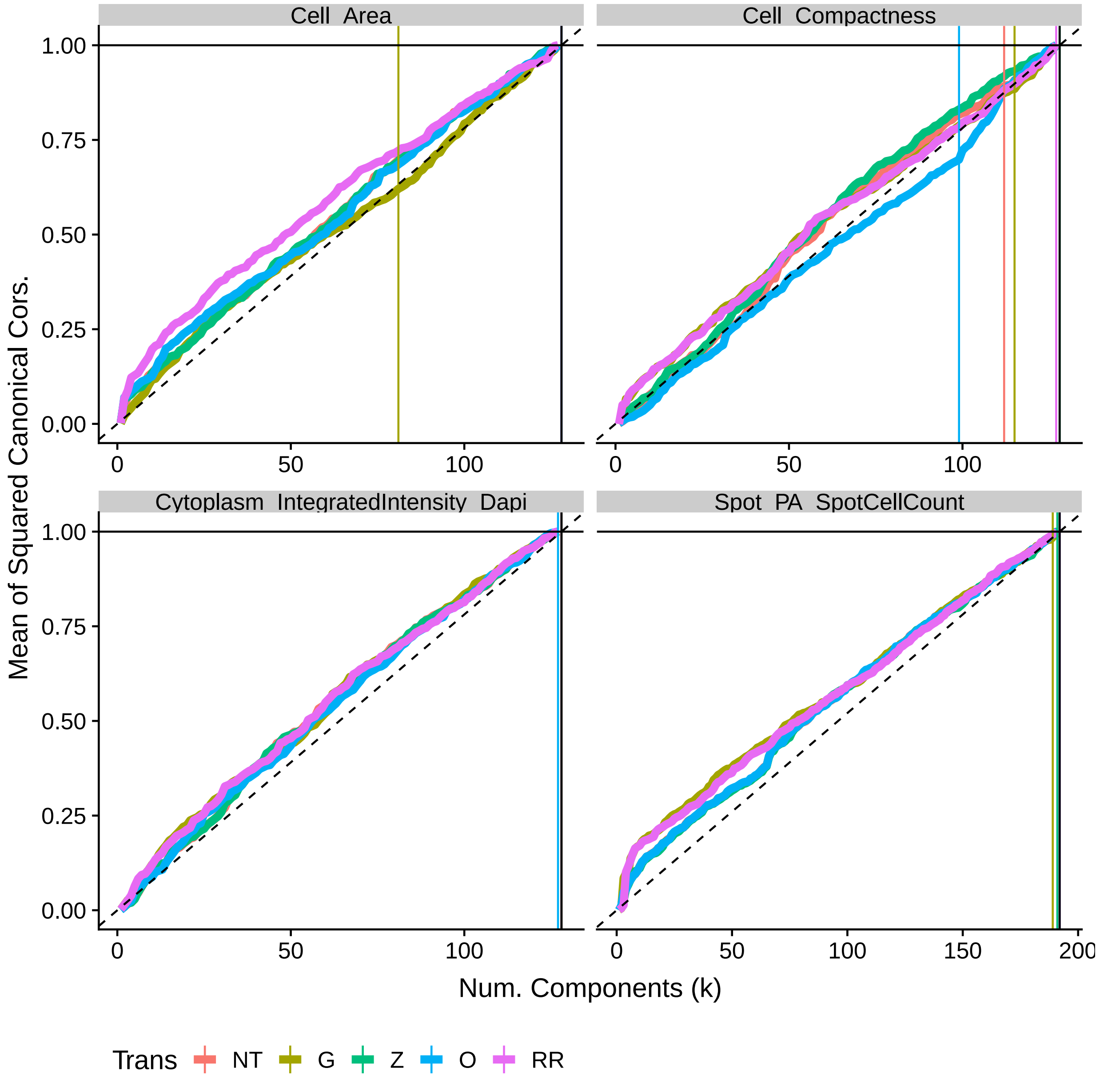


Figure 17: Similar to Figure 15 except correlation with well batch indicators.

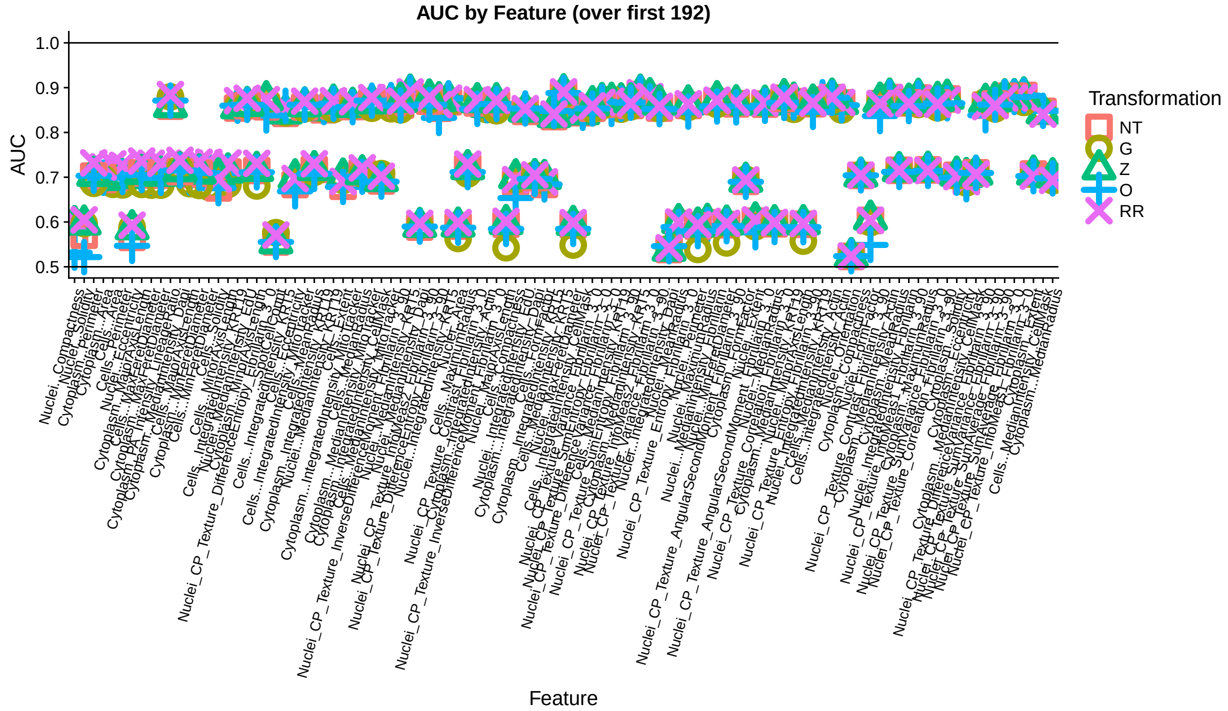


Figure 18: Similar to Figure 16 except correlation with well batch indicators.

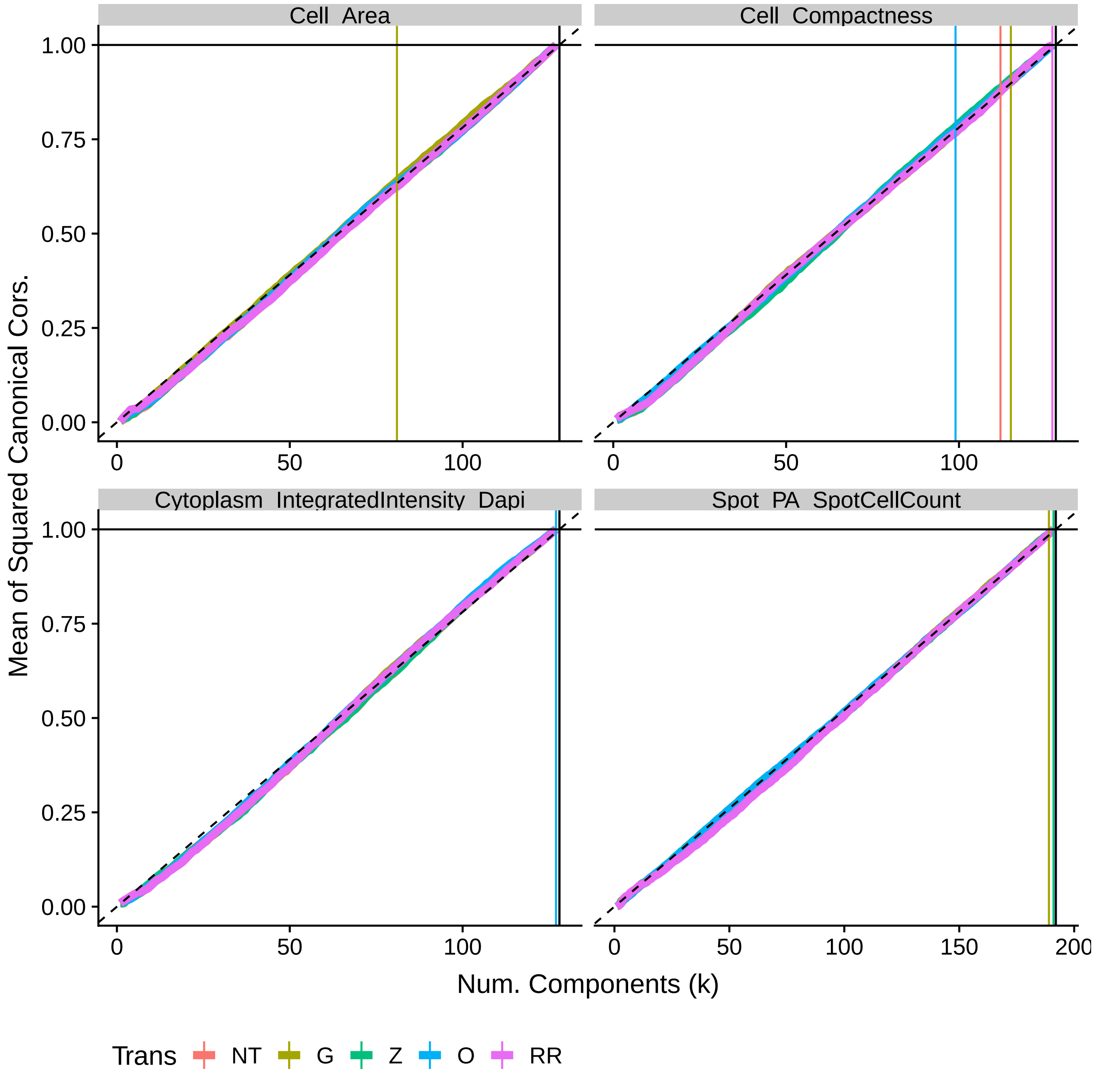


Figure 19: Similar to Figure 15 except correlation with ligand batch indicators.

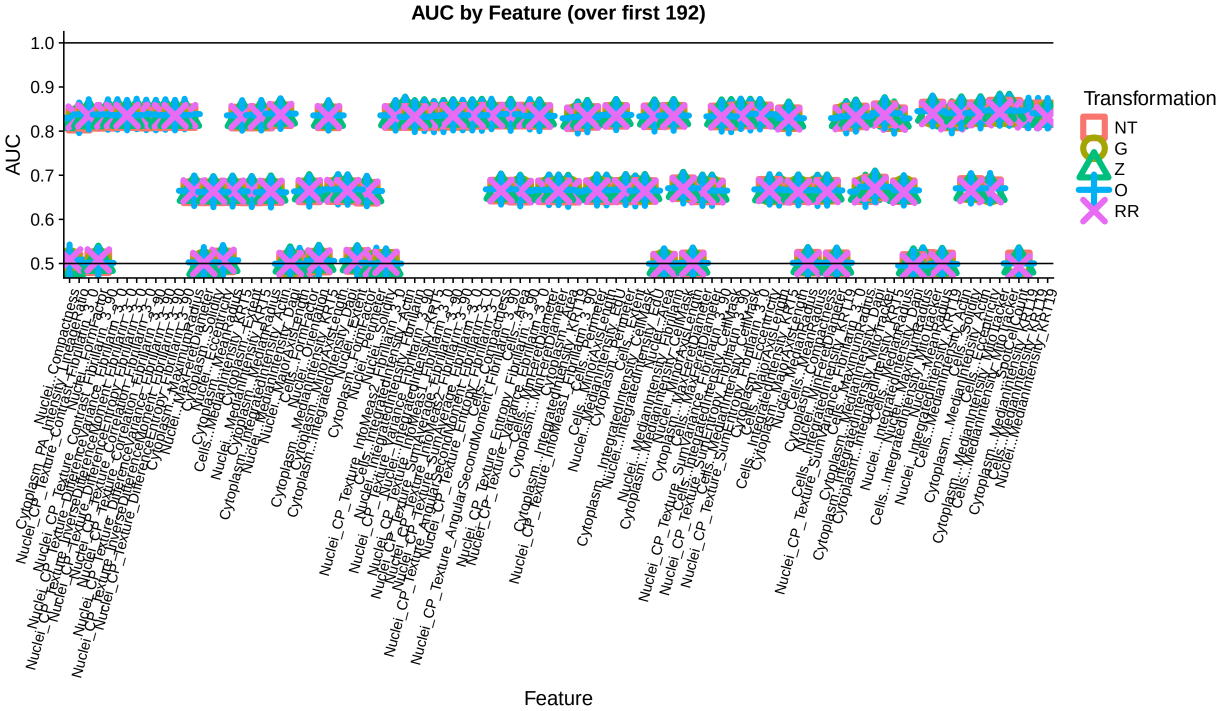


Figure 20: Similar to Figure 16 except correlation with well batch indicators.

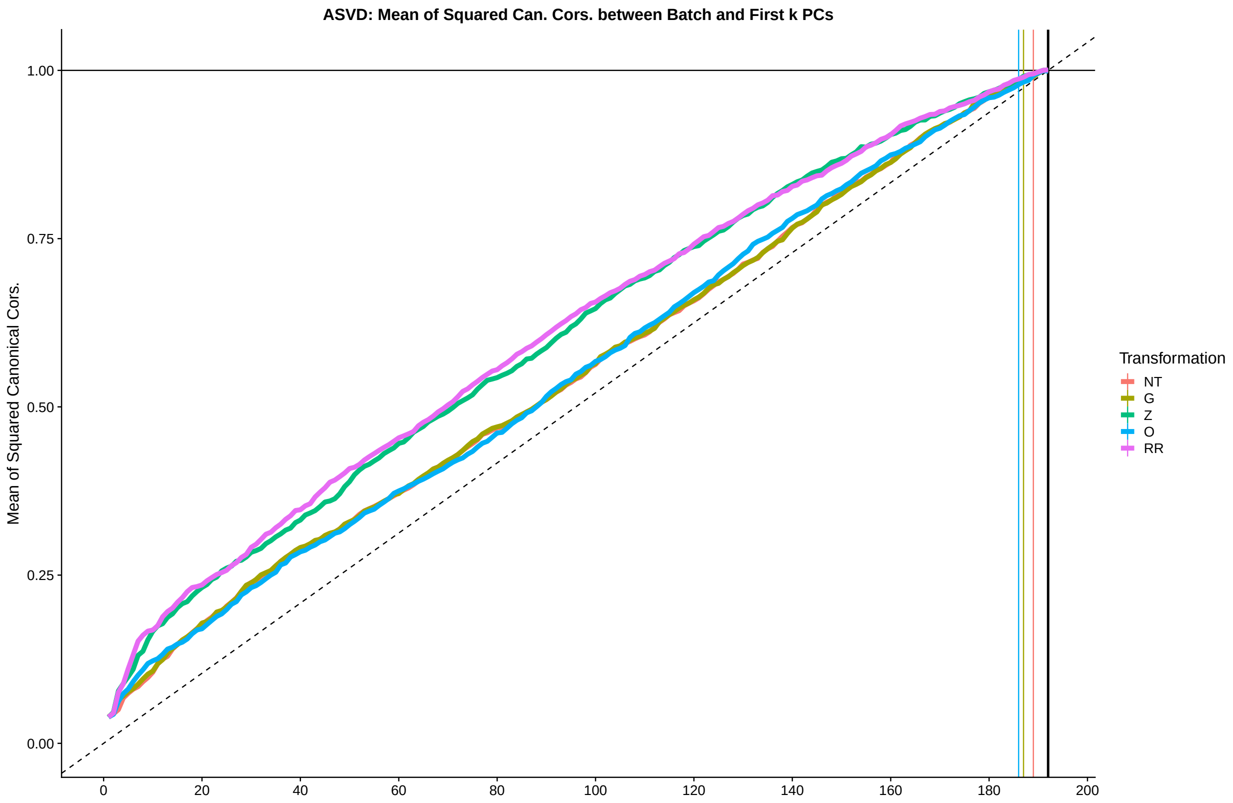


Figure 21: Mean of the squared canonical correlations between the first $k$ principal components and the plate indicator variables. Principal components come from integration of the 21 features that are measured across all MEMAs.

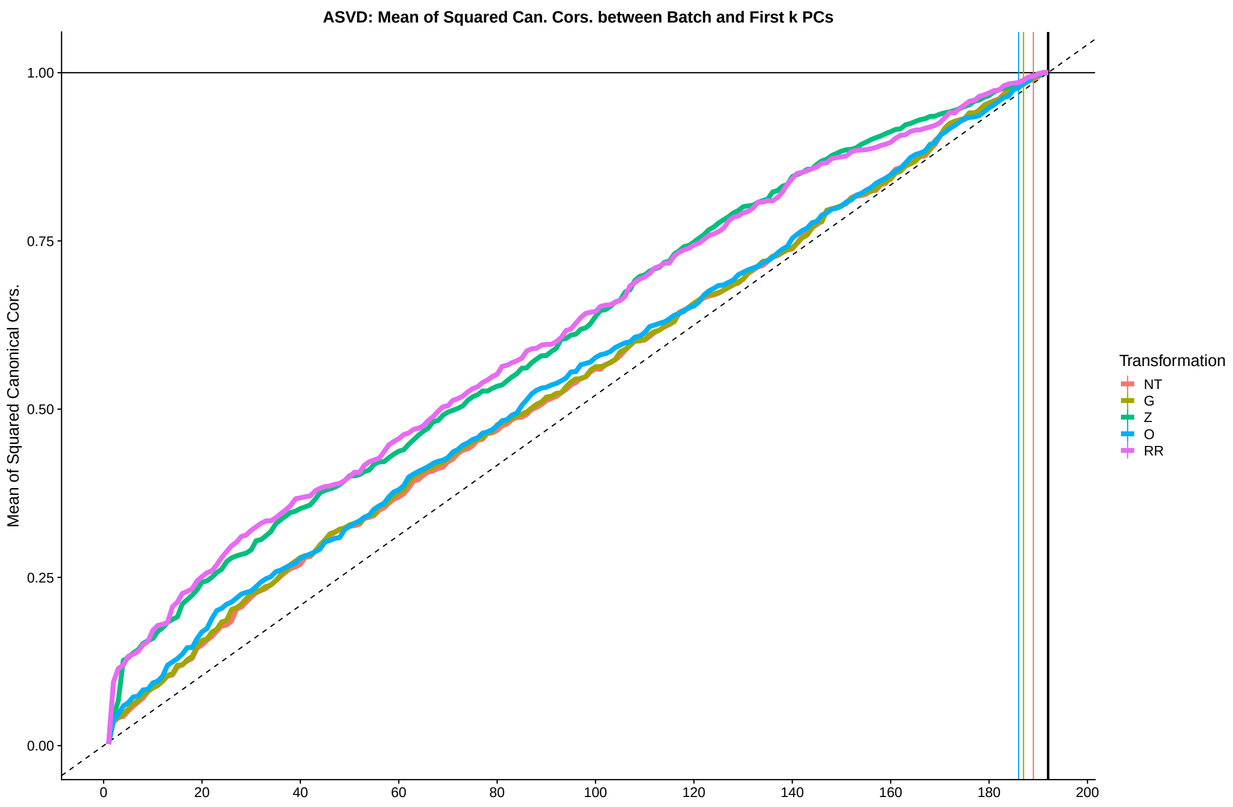


Figure 22: Similar to Figure 21 but calculating correlation with well indicators.

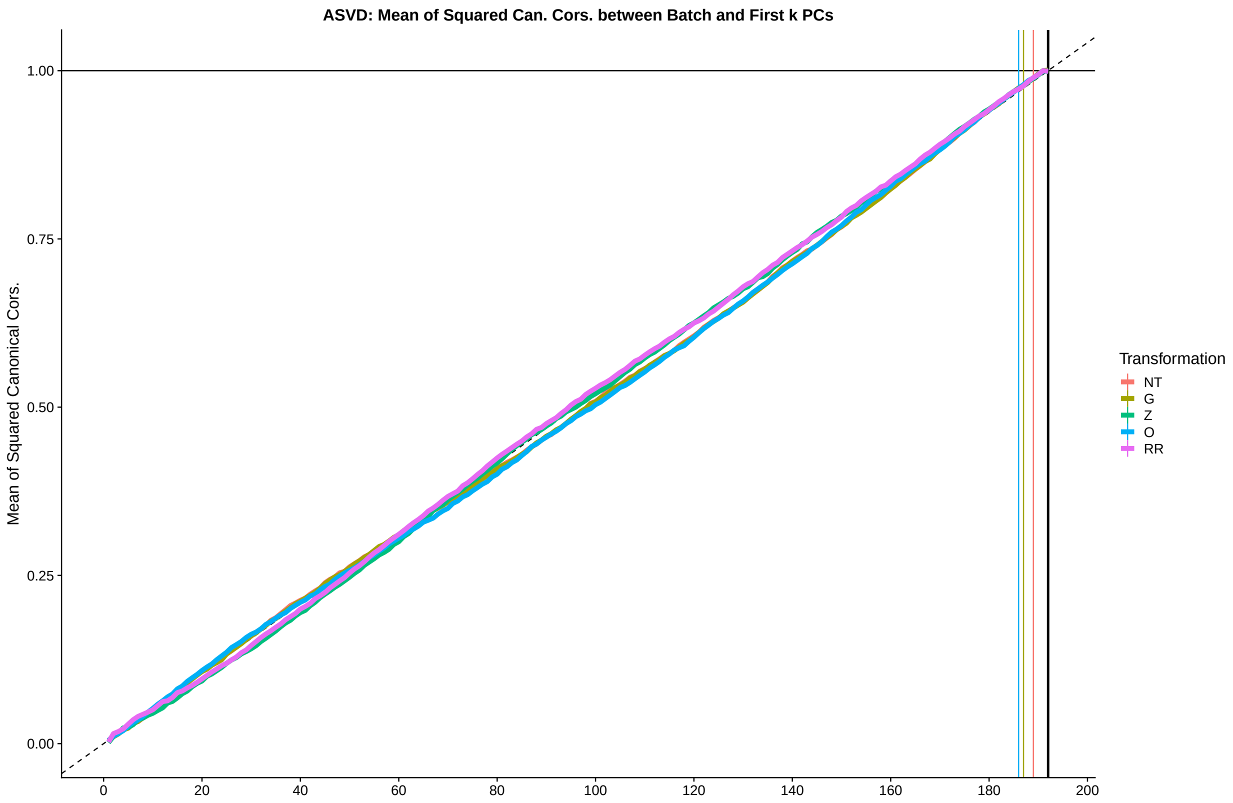


Figure 23: Similar to Figure 21 but calculating correlation with ligand indicators.

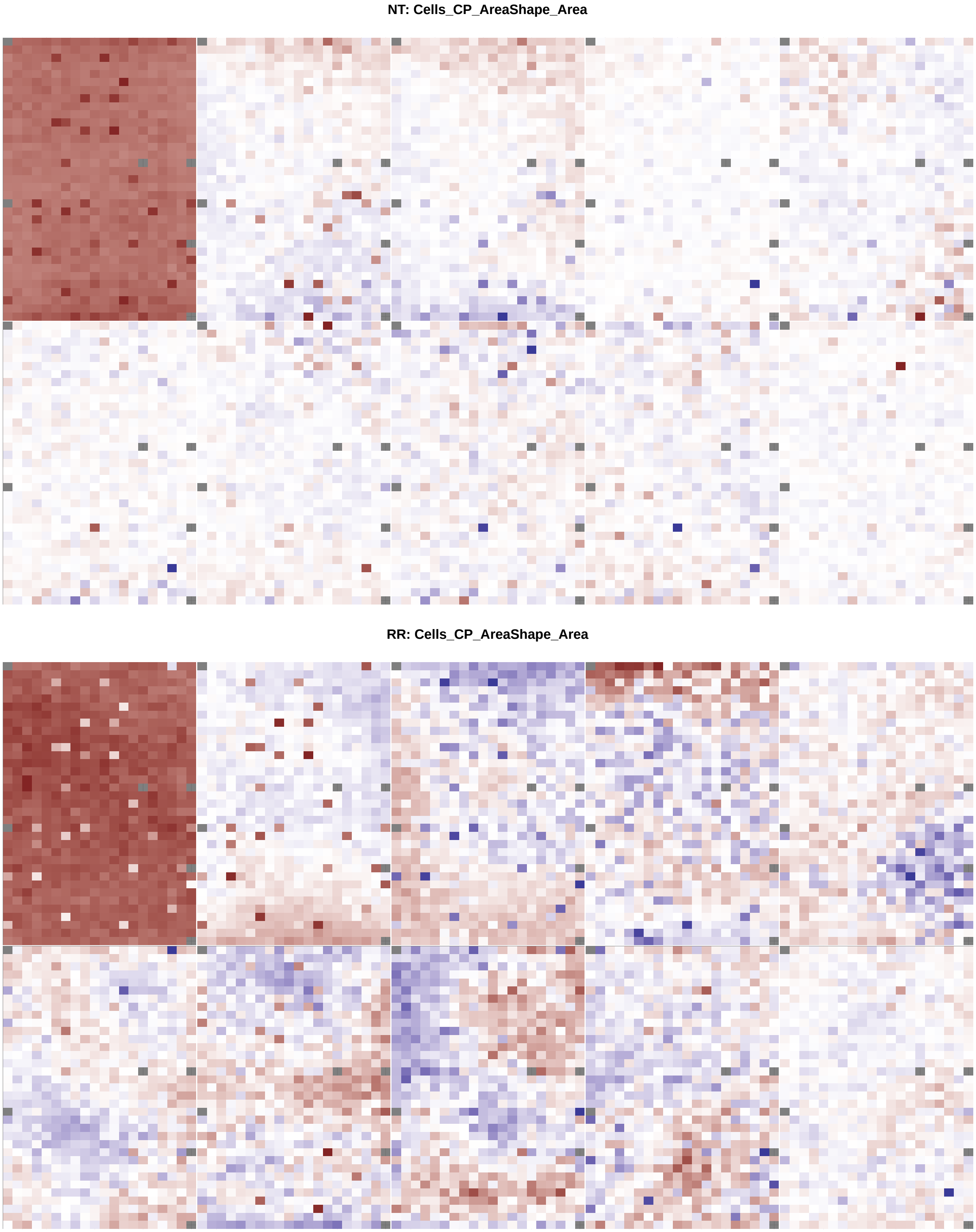


Figure 24: Heat map of elements of top 3 right singular vectors for the cell area feature.

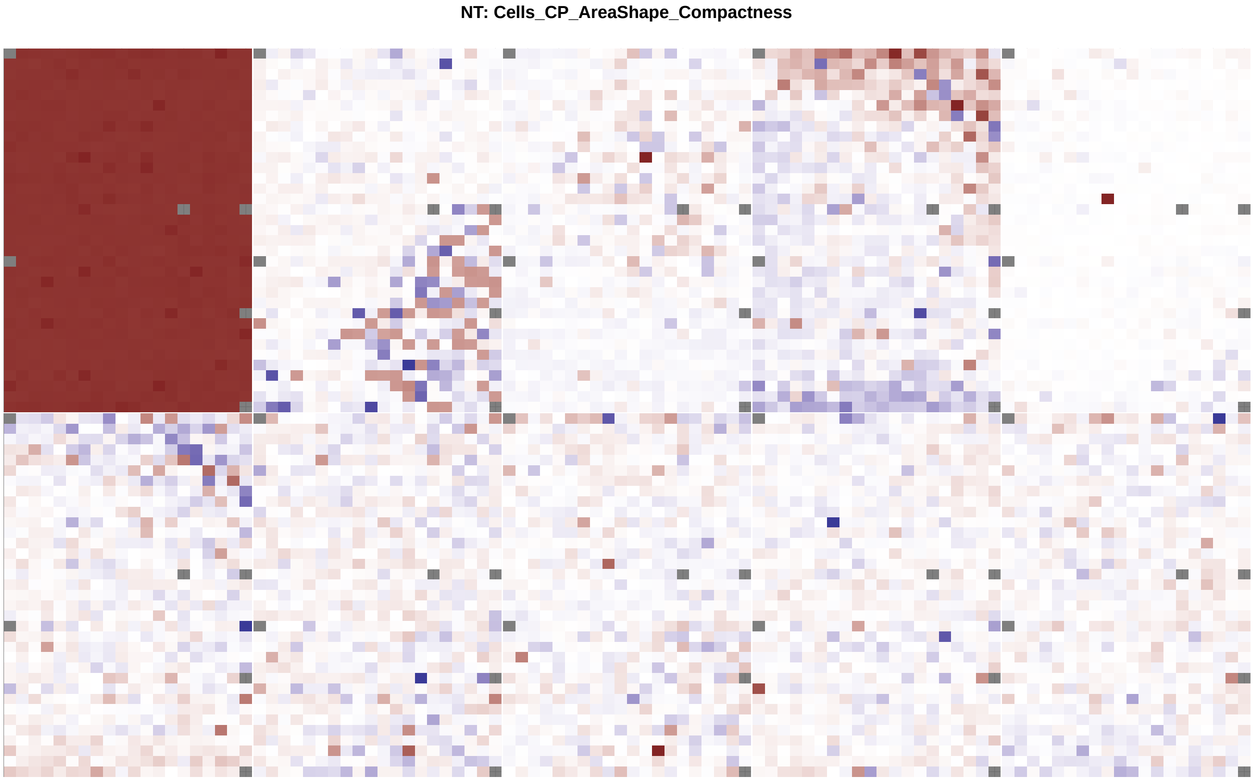


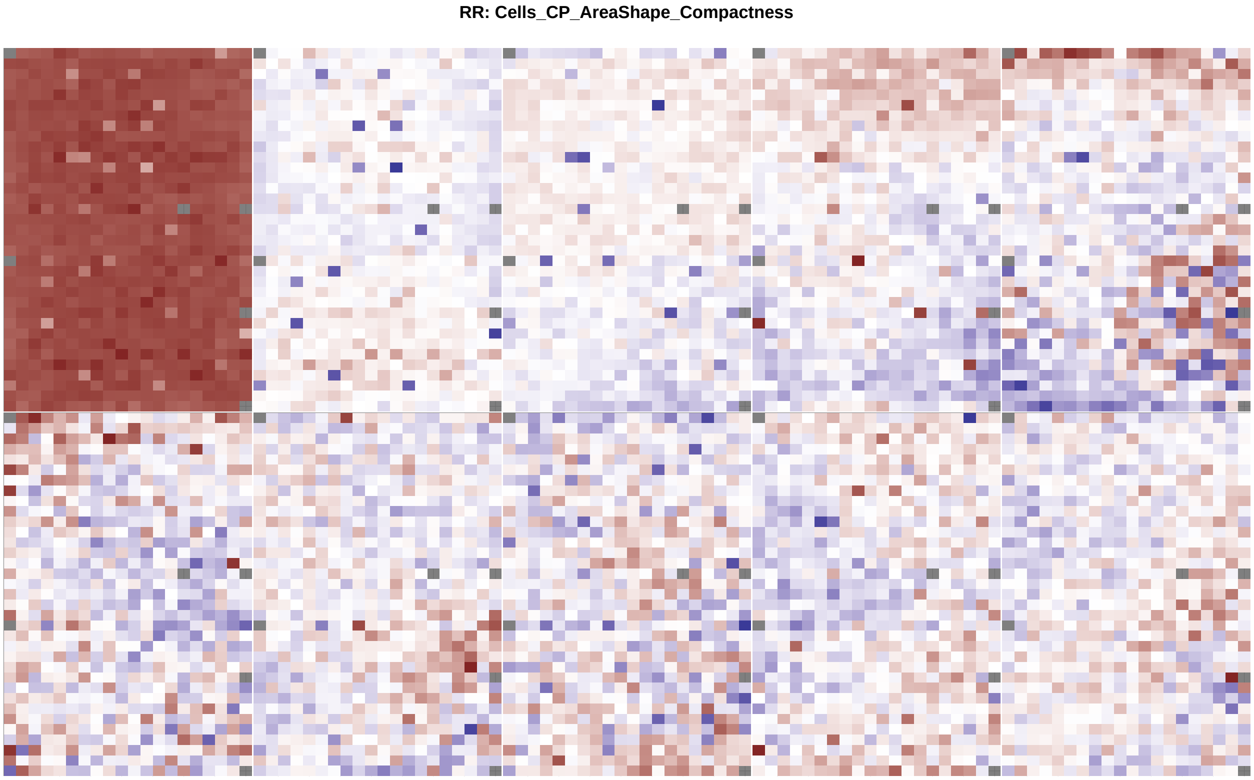


Figure 25: Similar to Figure 24 but for cell compactness feature.

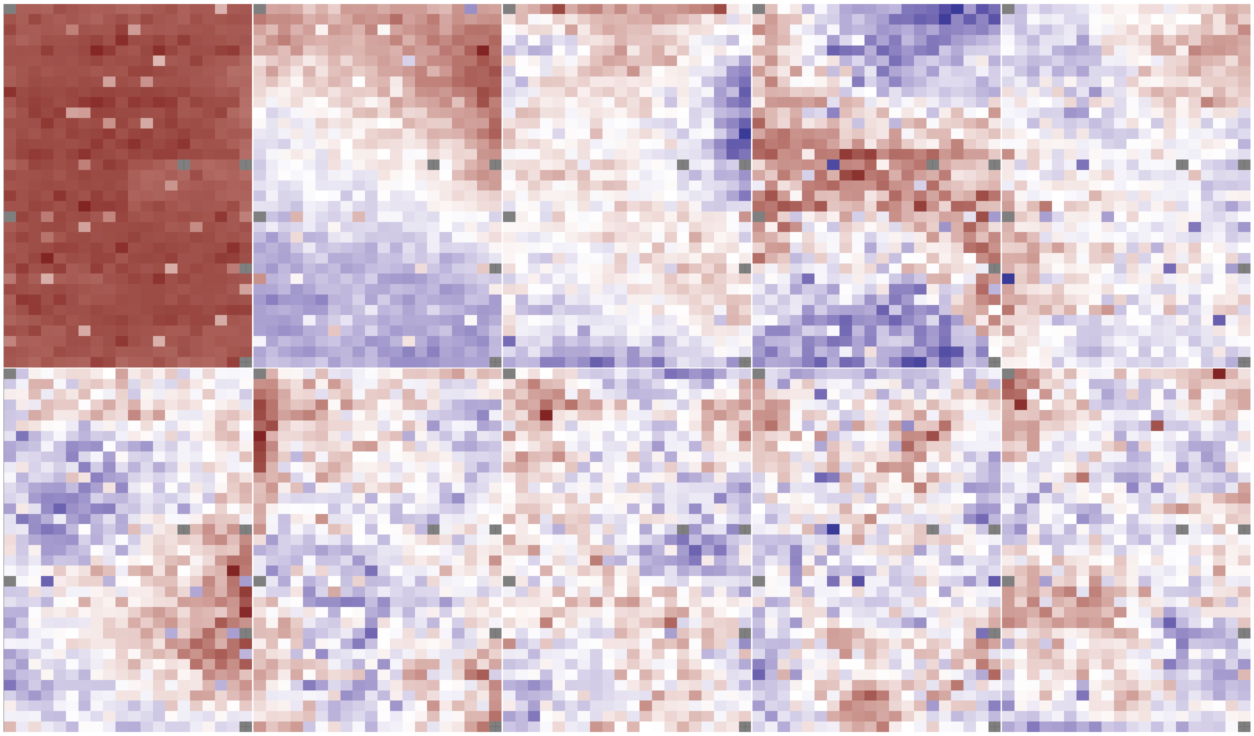


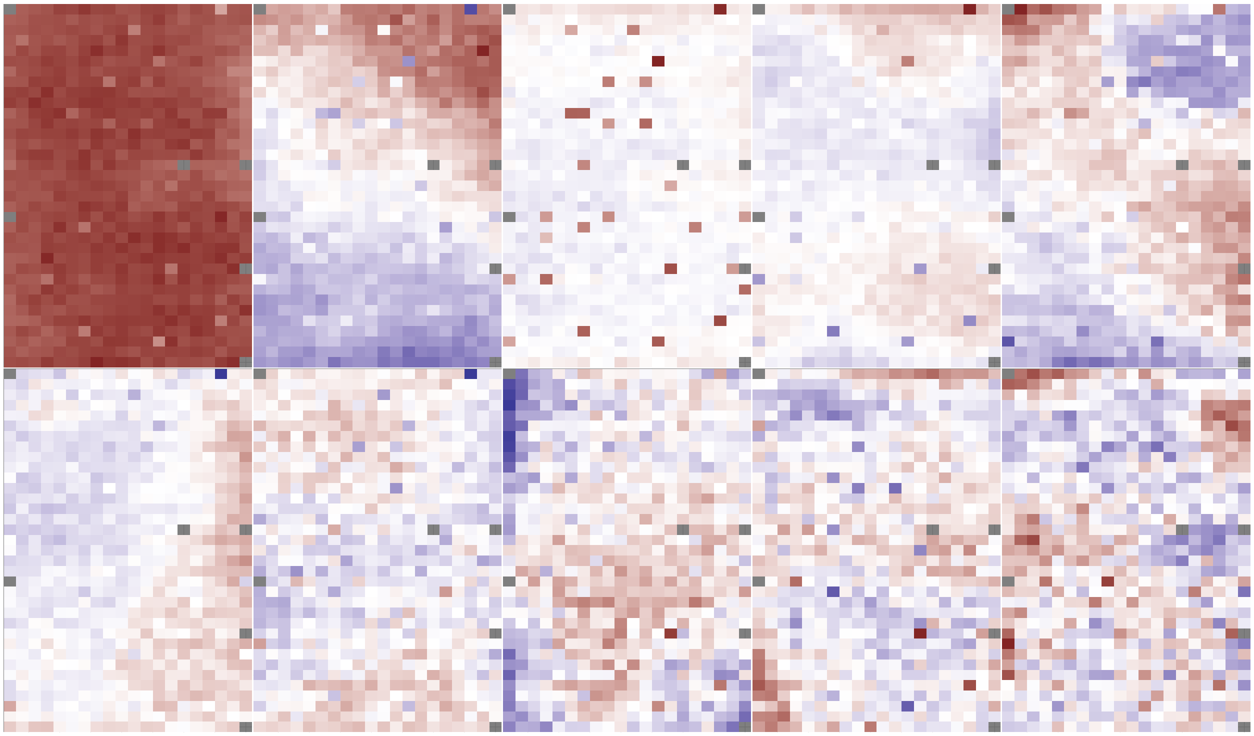


Figure 26: Similar to Figure 24 but for cell count feature.

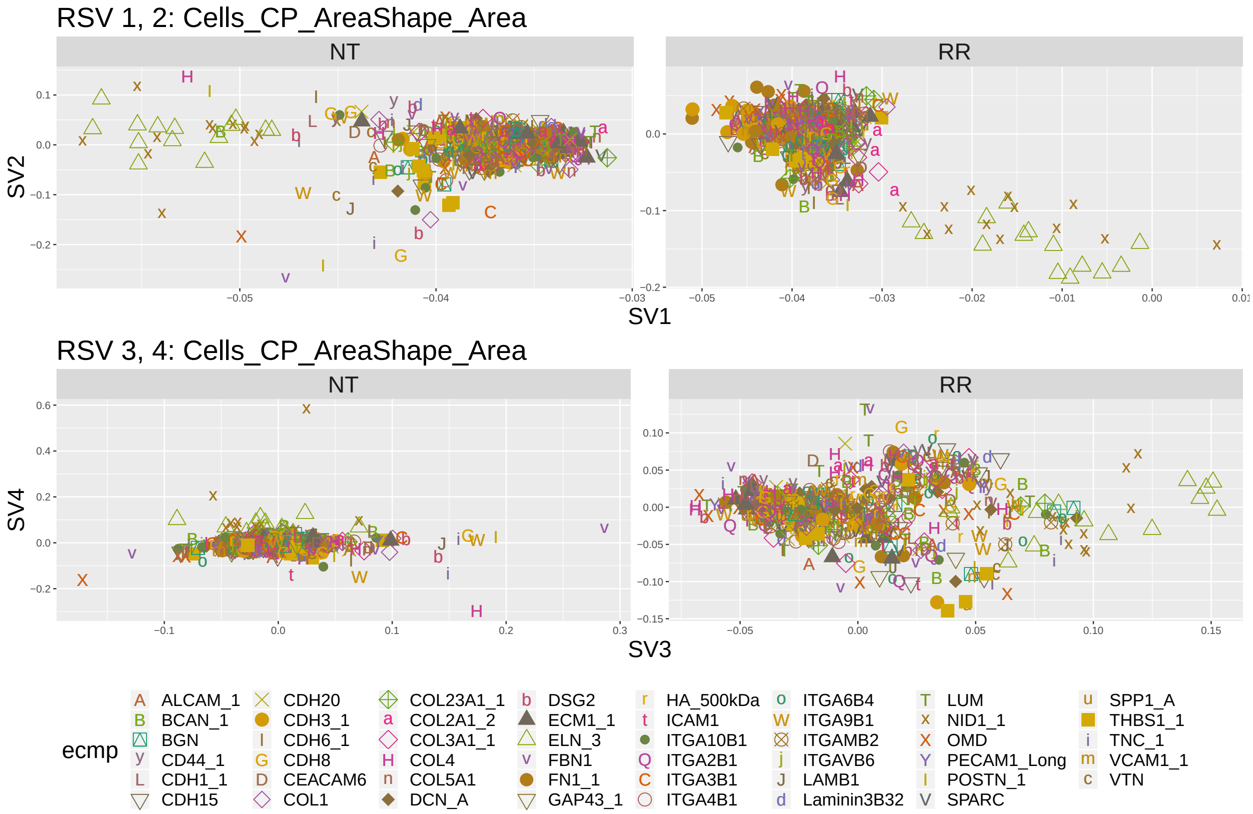


Figure 27: Scatter plot of elements of top two right singular vectors against each other for the cell area feature. Shape and color indicate ECMp of the spot corresponding to the elements of the singular vector.

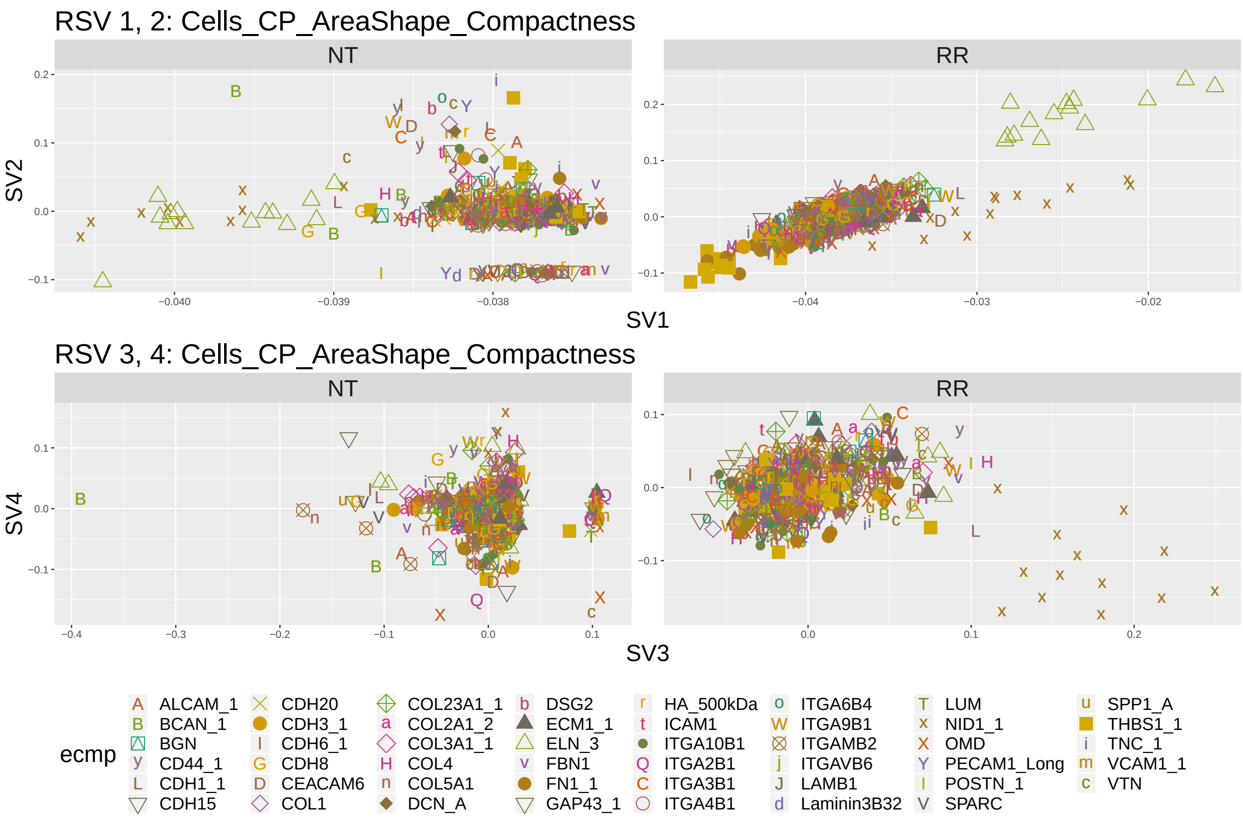


Figure 28: Similar to Figure 27 but for cell compactness feature.

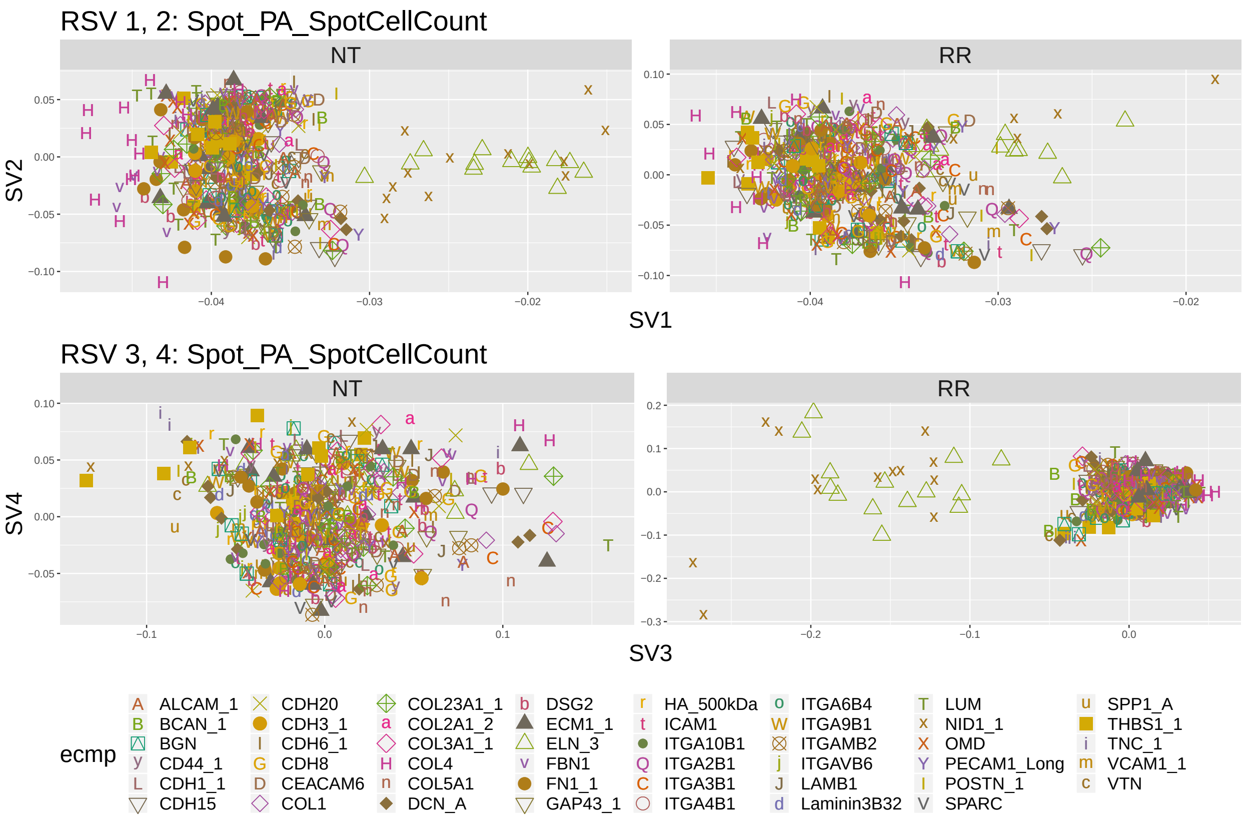


Figure 29: Similar to Figure 27 but for cell count feature.

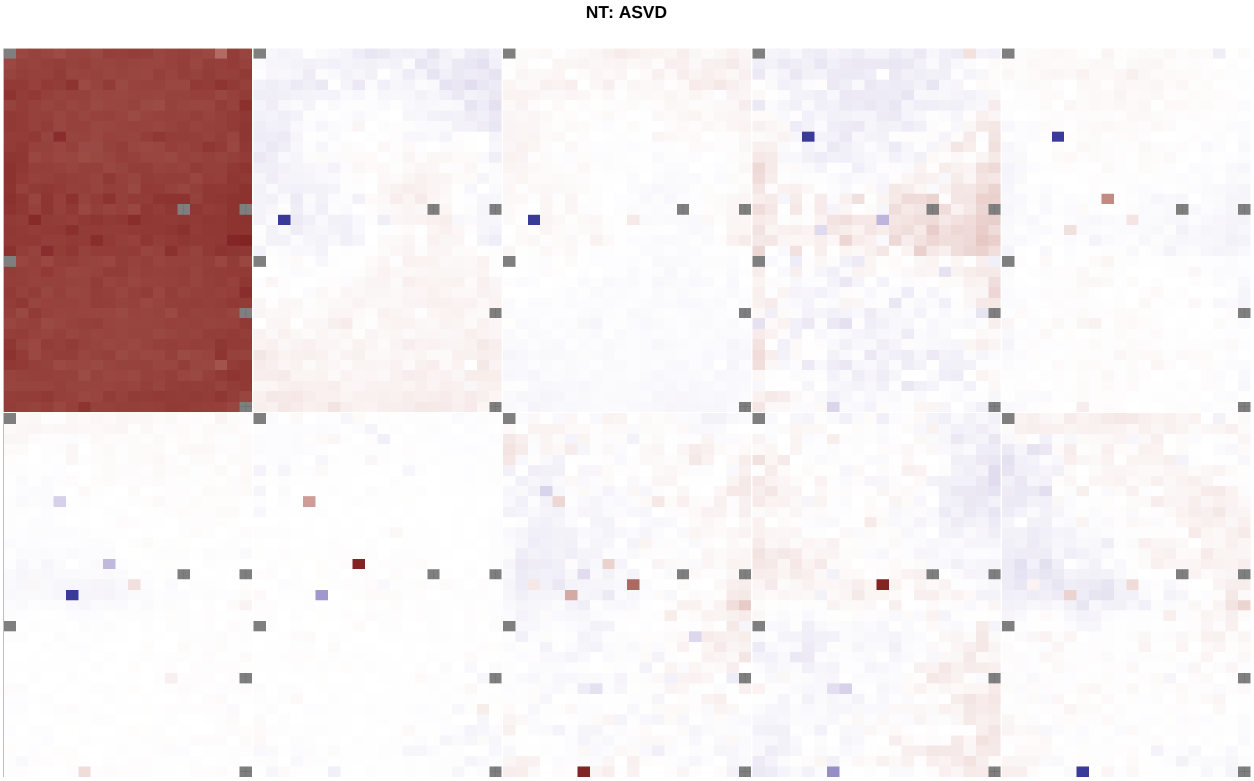


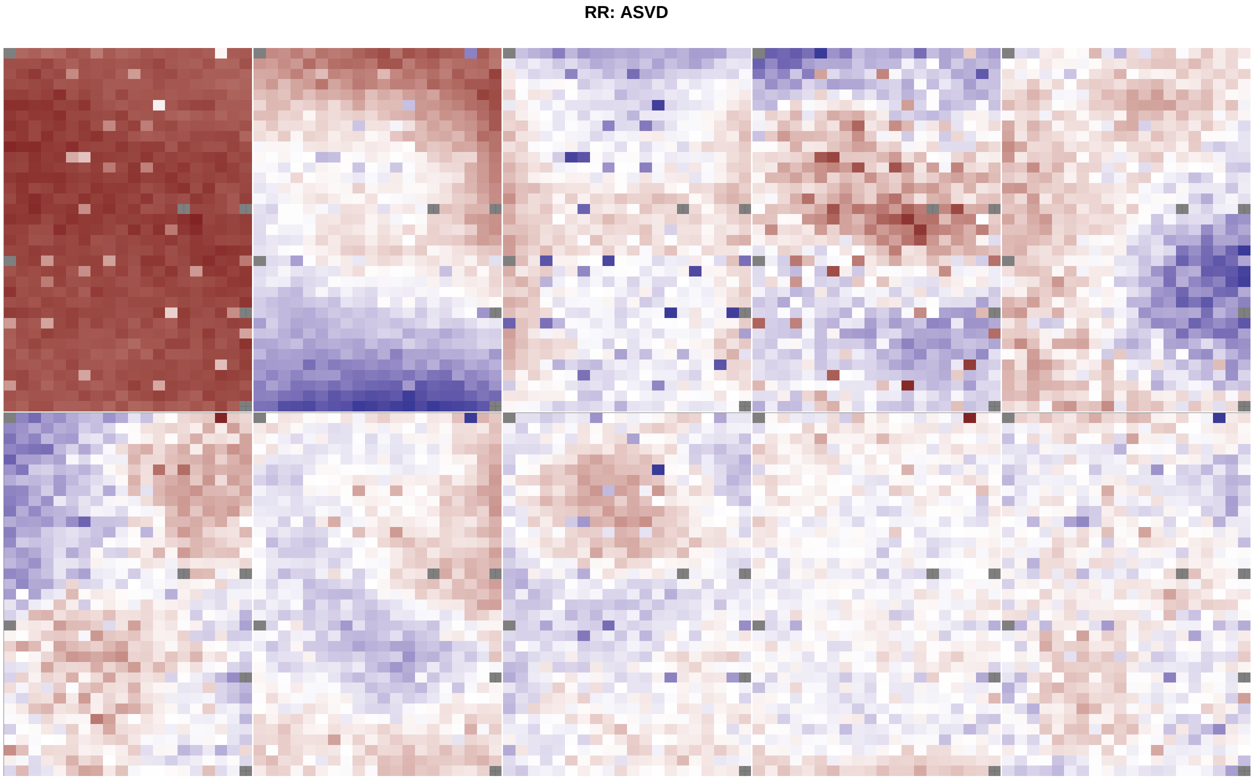


Figure 30: Heat-map of top 3 right ASVs calculated over 21 features measured on all MEMAs.

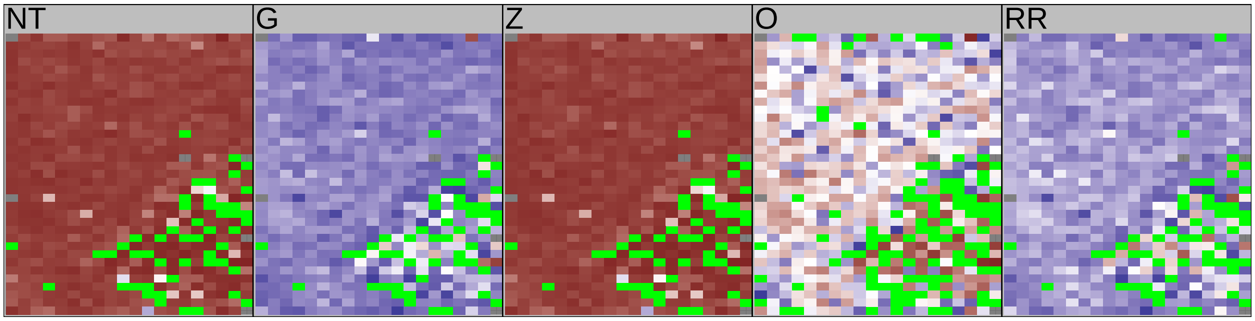


Figure 31: Missing values for well A03 on outlier plate LI8X00515. This plate is an outlier because it was processed using a different version of imaging processing software. Missing spots are indicated in green. Other colors indicate cell compactness feature. Notice that the missing values for (NT) are nearly identical to the the dark red spots in the second right singular vector in Supplementary Figure 25. This is what forms the group structure in Figure 11 as these missing spots are picked up on the outlying plate.
